# Human neonatal B cell immunity differs from the adult version by conserved Ig repertoires and rapid, but transient response dynamics

**DOI:** 10.1101/2020.08.11.245985

**Authors:** Bettina Budeus, Artur Kibler, Martina Brauser, Ekaterina Homp, Kevin Bronischewski, J. Alexander Ross, Andre Görgens, Marc A. Weniger, Josefine Dunst, Taras Kreslavsky, Symone Vitoriano da Conceição Castro, Florian Murke, Christopher C. Oakes, Peter Rusch, Dimitrios Andrikos, Peter Kern, Angela Köninger, Monika Lindemann, Patricia Johansson, Wiebke Hansen, Anna-Carin Lundell, Anna Rudin, Jan Dürig, Bernd Giebel, Daniel Hoffmann, Ralf Küppers, Marc Seifert

**Author notes:** contributed equally (Bettina Budeus, Artur Kibler, Martina Brauser, Ekaterina Homp). Correspondence: PD Dr. rer. nat. Marc Seifert, Institute of Cell Biology (Cancer Research), University Hospital Essen, Virchowstraße 173, 45147 Essen.

## Abstract

The human infant B cell system is considered premature or impaired. Here we show that most cord blood B cells are mature and functional as seen in adults, albeit with distinct transcriptional programs providing accelerated responsiveness to T cell-independent and T cell-dependent stimulation and facilitated IgA class switching. Stimulation drives extensive differentiation into antibody-secreting cells, thereby presumably limiting memory B cell formation. The neonatal Ig-repertoire is highly variable, but conserved, showing recurrent B cell receptor (BCR) clonotypes frequently shared between neonates. Our study demonstrates that cord blood B cells are not impaired but differ from their adult counterpart in a conserved BCR repertoire and rapid but transient response dynamics. These properties may account for the sensitivity of neonates to infections and limited effectivity of vaccination strategies. Humanized mice suggest that the distinctness of cord blood versus adult B cells is already reflected by the developmental program of hematopoietic precursors, arguing for a layered B-1/B-2 lineage system as in mice. Still, our findings reveal overall limited comparability of human cord blood B cells and murine B-1 cells.

**Significance Statement:** Neonates and infants suffer from enhanced susceptibility to infections. Our study contrasts with the current concept of a premature or impaired B cell system in neonates, by showing that most cord blood B cells are mature and functional. However, their responses are rapid but provide only short-term protection, which may help to improve infant vaccination strategies. We propose an altered perspective on the early human B cell system, which looks similar to but functions differently from the adult counterpart. Finally, our analysis indicates that cord blood- and adult B cell development occur layered as in mice, but certain mouse models still may offer a limited view on human neonatal B cell immunity.

## Introduction

Neonates and infants suffer from enhanced susceptibility to infections (1) and four out of ten childhood deaths worldwide are due to infectious diseases (2, 3). Birth is a dramatic transition for the neonate, exposing it to an environment with a high microbial burden compared to that *in utero*, and the infant immune system itself is considered immature to meet these challenges (1, 3, 4). Low neutralizing antibody titres are a critical limitation in infant immunity, and current vaccination strategies aim at generating protective seroconversion (3, 5). T cell-independent (TI) polysaccharide vaccines either fail (6) or elicit short-lived responses with quickly waning IgG titres (7) and do not induce B cell memory (8), but rather hyperresponsiveness (immune exhaustion) (9). Similarly, T cell-dependent (TD) conjugate vaccinations are generally of low magnitude and do not persist well in early infancy (5, 10). Immature infant B cell immunity is associated with imbalances in T cell support (4, 11), and further evident by the importance of maternal-derived antibodies in neonatal immunity (1). Several reasons for infant B cell immaturity have been described.

Compared to adult naïve B cells, human neonatal-derived B cells (“neonatal B cells”) were reported to express lower levels of important surface molecules, including CD21 (12), TNFRSF13B (13), CD73 (14), CD22 (15), CD62L and CCR7 (16), CD40, CD80, and CD86 (13), all of them associated with a lower magnitude of B cell responses. Differences in TLR expression influence innate and adaptive immunity of human neonates (17) and B cell receptor (BCR) signal transduction may differ significantly from adult-derived B cells (“adult B cells”), depending on the stimulus (15, 18). In neonatal B cells, IgV gene mutations are infrequent (19–21), affinity maturation is delayed (3), and IgG/IgA class switching is impaired (18), even upon co-stimulation with IL-21 (17).

Multiple immune factors contribute to the lower magnitude of infant responses to vaccination and enhanced vulnerability to infections compared to adults (3, 5). In mice, the interaction of neonatal lymphocytes and dendritic cells (DC) is immature, resulting in delayed maturation of follicular DC networks (22). Human infants also show limited germinal centre (GC) reactions (23, 24). Newly generated plasma cells (PC) compete for limited access to survival niches (25–27), and limited survival signals (28). Finally, single vaccine doses at birth can fail to elicit seroconversion while priming for secondary responses (5). The low IgV gene mutation load in neonatal B cells was proposed to favour memory B cell over PC differentiation (3, 29). In light of these multiple limitations of infant immunity, B cell-intrinsic differences are considered unlikely by themselves to provide an insurmountable obstacle to strong responses (1).

Current knowledge of the infant B cell system is mostly derived from vaccination studies or is inferred from mouse models. In mice, neonatal B cells include developmentally and functionally distinct B cell lineage (B-1 cells)(30) whereas adult haematopoiesis produces conventional B-2 cells (31). Among other markers, CD5 is used to distinguish murine B-1a (CD5^+^) from B-1b (CD5^-^) cells. B-1 and B-2 cell transcriptome patterns diverge (32–34). BCR specificity and signalling strength influence fetal B cell development (35) and can instruct B-1 phenotypes in adult B cells (36). Fetal, but not adult B-1 cell Ig repertoires show a proximal-biased V-gene usage (37), and limited addition of non-germline-encoded (N) nucleotides (38, 39). Neonate mice exhibit pre-immune clonal B-1 cell expansions (40). B-1 cells display a primitive, innate-like functionality and produce natural antibodies, independent of T cell help (41). In humans, the existence of a B-1 cell counterpart is debated (19, 20, 42–50). CD5 expression in humans is not confined to a fetal B cell lineage (51–54) and human neonatal B cells share only a few of the defects observed in mice (1, 16).

Our understanding of human infant B cell immunity is limited. The current view implies neonatal B cell functional immaturity in responsiveness and differentiation, particularly under TD stimulation, and their interaction with other immune cells is considered less developed.

For a comprehensive characterization of human neonatal mature B cells, we analysed cord blood B cell Ig repertoires, molecular patterns, and functional capacities. We show here that cord blood B cells are phenotypically similar but highly distinct in development and function from adult B cells. In contrast to current belief, the vast majority of cord blood B cells consists of mature and fully functional B lymphocytes that respond faster and more efficient to autologous T cells, but also innate stimulation compared to adult naïve B cells. Cord blood B cells show strong IgA switching capacity and their differentiation potential is mostly confined to antibody-secreting cell (ASC) fate. Their molecular and functional distinctness is supported by a conserved cord blood Ig repertoire as up to 8% of BCR clonotypes among 180,000 BCR rearrangements analysed are identical between individual neonates. Finally, we show both similarities and differences between human cord blood B cells and murine B-1 cells, suggesting limited comparability of their respective molecular patterns between both species.

## Results

### A Major Fraction of Human Umbilical Cord Blood B Cells is Mature

Human neonates were reported to have a high frequency of CD5^+^CD10^+^CD24^+^CD38^high^ transitional B cells with immature functionality (53, 55). To investigate whether human neonatal B cell responses are limited by mature B cell numbers, we determined the average number of mature and transitional B cells in a cohort of 43 umbilical cord blood (UCB) samples. Our analysis showed that on average, neonates have similar proportions of B cells among lymphocytes as adults (10% of mononuclear cells (range 3% to 18%), compared to 11% (range 4% to 20%) in peripheral blood (PB) of adults > 20 years). Indeed, the fraction of transitional B cells (CD5^+^CD10^+^CD24^+^CD38^high^) is higher in cord blood than in adult PB. However, most cord blood B cells (64%, range 48% to 79%) shows a mature naïve phenotype (CD19^+^CD10^-^CD24^-^CD27^-^CD38^low^ IgM^+^IgD^+^) (Figures 1A and B and S1A and B). Our initial staining panel indicates that the B cell pool in cord blood includes a major fraction of IgM expressing (likely foreign antigen-naïve) mature B cells. To allow for comparison of age-related changes (and to assess similarities to murine B-1a and B-1b cells) we analysed mature CD5^-^ (CD5^-^CD23^+^CD27^-^CD38^low^IgD^+^) and CD5^+^ (CD5^+^CD23^+^CD27^-^CD38^low^IgD^+^) B cells derived from human cord blood and adult PB separately. On average cord blood mature B cells are composed of equal-sized fractions of mature CD5^-^ and CD5^+^ B cells (Figures 1A and S1C), the CD5^+^ fraction decreases with age (Figure 1C). Notably, cord blood B cells express higher levels of surface IgM compared to adults naïve B cells (Figures 1A and S1A).

**Figure 1.**
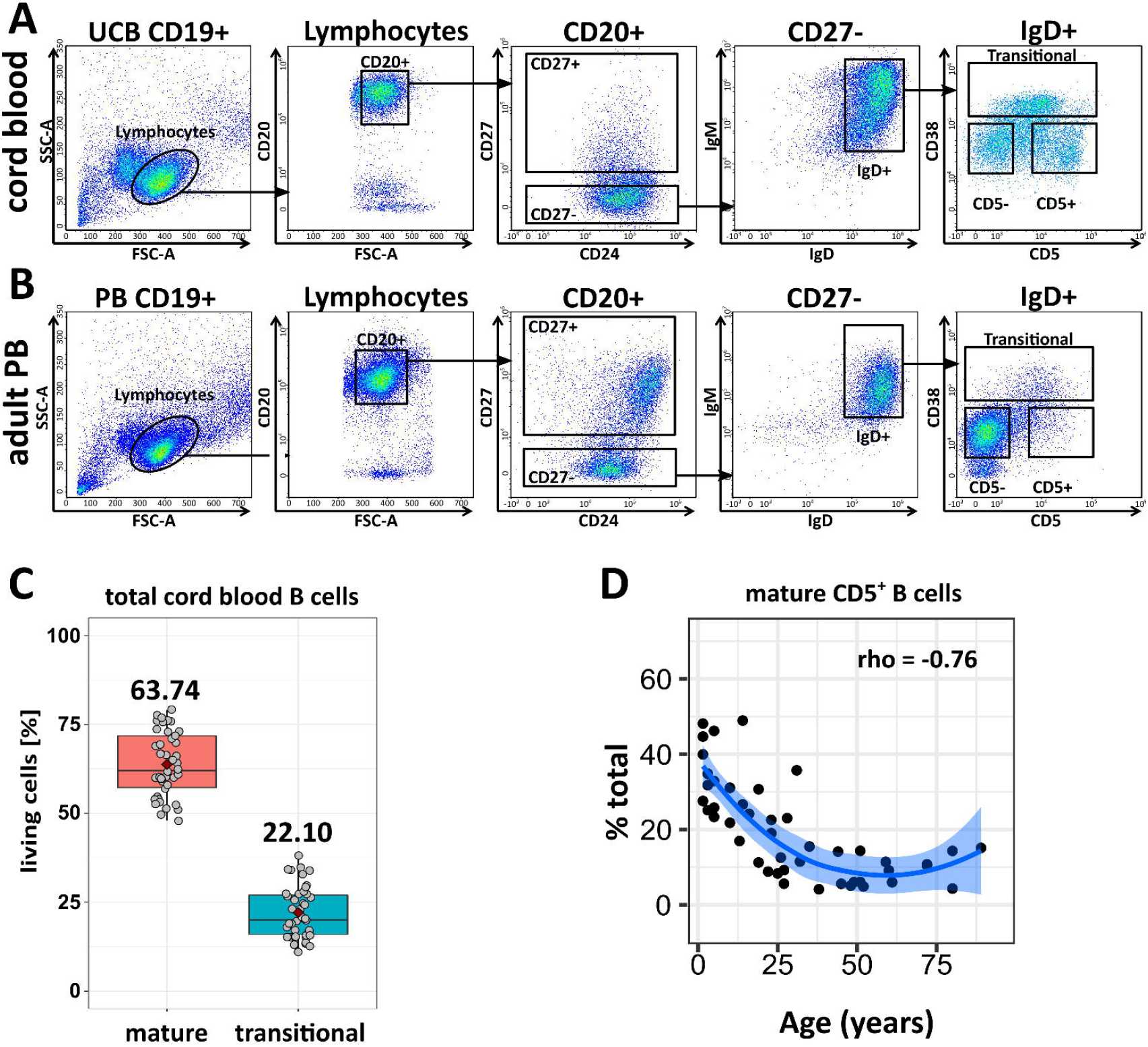
Quantification of Human Cord Blood and Adult Mature B Cells. (A) Gating strategy for cord blood and adult mature B cell subsets. (B) Proportions of mature (CD5^+^ and CD5^-^) B cells and transitional B cells in 43 randomly selected neonates (UCB). (C) Age-dependent changes in the proportion of mature CD5^+^ B cells from 45 donors aged 0 – 90 years. Line of fit (bold line) and 95% confidence interval (shaded area) were calculated by R smooth.

We further quantified memory B cells (CD27^+^), plasmablasts (CD27^high^CD43^+^) and the recently proposed CD27^+^CD43^+^ human B-1 cells (Figures 1A and S1D) (44), according to their of surface IgM and IgD expression (Figure 1A and data not shown). Mature cord blood B cells barely include memory B cells and plasmablasts, or CD27^+^CD43^+^ B cells whose numbers increase with age until adulthood, (Figure S1D), in line with previous publications (55, 56). We conclude that a major fraction of human umbilical cord blood B cells is mature.

### Cord Blood and Adult Mature B Cells have Distinct Transcriptomes

Despite the presence of a large fraction of B cells with a mature phenotype in cord blood, neonates and infants have an elevated susceptibility to infections compared to adults, which could be attributed to functional impairment of the B cell system.

Transcriptome-wide comparison is a reasonable approach to estimate functional differences between B cell subsets (52, 57). To this end, we generated RNAseq profiles of sort-purified (Figure S1B) CD5^-^ (CD5^-^CD23^+^CD27^-^CD38^low^IgD^+^) and CD5^+^ (CD5^+^CD23^+^CD27^-^CD38^low^IgD^+^) mature B cells from cord blood, and adult naive CD5^-^ (CD5^-^CD23^+^CD27^-^CD38^low^IgD^+^) and CD5^+^ (CD5^+^CD23^+^CD27^-^CD38^low^IgD^+^) mature B cells from PB for comparison. Human adult IgM memory (CD5^-^CD23^-^CD27^+^CD38^low^IgM^+^IgD^+^) and IgA or IgG class switched memory (CD5^-^CD23^-^CD27^+^CD38^low^IgG^+^ or IgA^+^) B cells from PB and tonsil served as analytical root for a robust determination of the relative similarity between adult and cord blood B cell subsets.

According to a t-SNE analysis of the top 10,000 expressed genes of each subset analysed (Figure 2A), and a hierarchical clustering of the 4,400 most variable transcripts (Figure 2B), human cord blood B cell transcriptomes are unique and deviate from adult B cell samples. Main components of the enriched transcription patterns represent “signalling by the BCR”, “MHC-I mediated antigen presentation” and “pre-mRNA processing” as determined by Reactome pathway enrichment analysis (Figure S2C). Gene set enrichment analysis (GSEA) revealed numerous surface receptors and signalling pathways enriched in cord blood versus adult B cells (Table S1). Enriched signatures included multiple gene sets associated with mitogen responses, BCR- and CD40 signalling (Figure 2C), suggesting that cord blood B cells are transcriptionally prepared to respond to TI-I, TI-II and TD stimulation, respectively. Moreover, TGFβ signalling pathway as prerequisite for IgA switching (58), and multiple gene sets associated with TNF-receptor-, interleukin-, and chemokine-signalling suggested that cord blood B cells have broad response capabilities in both TI and TD immunity (Table S1 and Figure 2C). The deviant transcriptional character of cord blood and adult B cells is not only reflected by coherent gene signatures, but also evident on single molecule level. Figure 2D shows a supervised selection of central molecules in BCR signalling, B cell activation or T cell-B cell-interaction, spanning up to four-fold change in expression. In depth analysis of the enriched pathways shows that single gene sets are not simply enriched, but may also show a distinct pattern of higher or lower expression of specific signalling components between cord blood versus adult B cell subsets (e.g. BCR signalling in cord blood: LYN and PLCG2 versus adult: SYK, BTK and BLNK).

**Figure 2:**
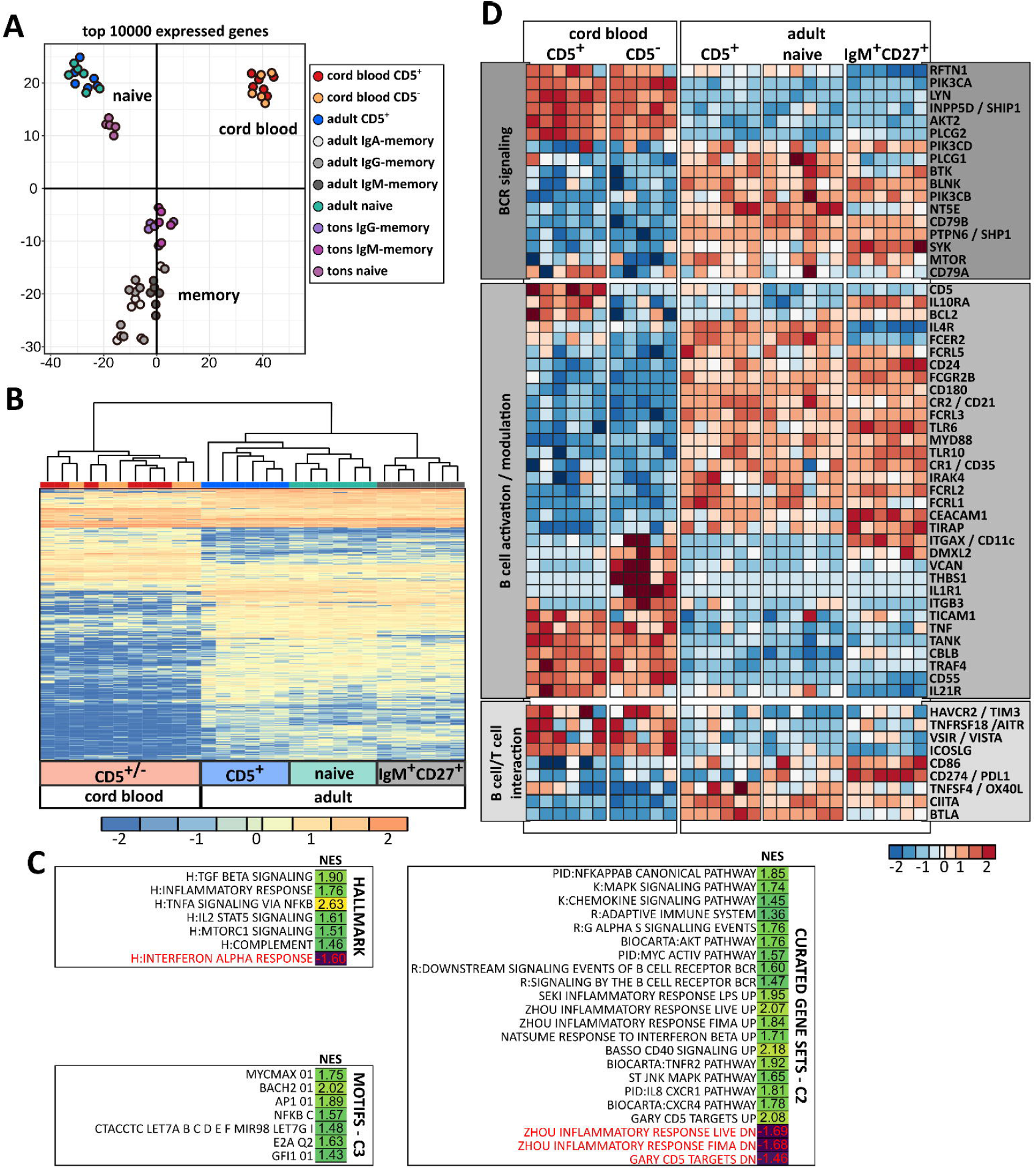
Transcriptome-wide Comparison of Human Cord Blood and Adult Mature B Cells. (A) T-SNE plot (R-package Rtsne and standard settings, https://CRAN.R-project.org/package=Rtsne) showing the distribution of different B cell subsets based on the top 10,000 expressed genes. (B) Hierarchical clustering and heat map showing the 4,400 most variable transcripts (MANOVA, q<0.05) of cord blood and adult PB mature CD5^+^ and CD5^-^ B cells, and adult PB IgM memory B cells. The colouring spans 4-fold differences, row-wise normalized (legend). (C) Gene set enrichment analysis of hallmark, C2, and C3 gene sets. Shown are gene sets enriched (black) or depleted (red) in cord blood versus adult B cells with a p-adjusted value below 0.05 with their normalized enrichment score (NES) (R package fgsea, doi: 10.1101/060012). This selection focusses on immune signalling, for the total list, see Table S1. (D) Detailed heatmap for selected genes involved in BCR signalling, B cell activation/modulation, B cell/T cell interaction, and B-1 cell development among cord blood and adult mature CD5^+^ and CD5^-^ B cells, and adult IgM memory B cells. The colouring depicts row-wise normalized fold-changes.

CD5 expression discriminates adult and cord blood subsets only marginally on transcriptional level, by a differential expression of 87 (out of 17,000) transcripts (Figures S2A and B). However, transcriptional differences between CD5^+^ and CD5^-^ subset are mostly mild, and their biological relevance is thus, questionable.

Arid3a and Bhlhe41 expression are characteristic for murine B-1 cells (59–61). Single cell analysis of *BHLHE41* expression by RNA flow cytometry revealed low levels in cord blood B cells, while higher expression was largely restricted to the adult memory compartment (Figure S2E). Moreover, the RNAseq profiles of cord blood B cells did not show a consistent enrichment of typical murine B-1 cell expression patterns (Figures S2D and F) (32–34).

We conclude that human cord blood B cells show unique transcriptome patterns, suggesting efficient responsiveness to both TI and TD stimulation, but barely share signatures with murine B-1 cells.

### Cord blood and Adult B Cells are Phenotypically Similar, but Respond Differently

Our transcriptome analysis supports the idea of many differentially expressed genes with high impact on B cell responsiveness. To validate differential expression, which does not directly translate to protein expression levels (62, 63), we selected 29 surface molecules and performed flow cytometric analysis on a total of 30 cord blood and 21 adult PB samples.

Flow cytometric analysis did neither reflect the transcriptional differences nor support the previously reported lower levels in T cell-B cell-interaction molecules or CD21 expression on resting cord blood B cells (Figure 3A and compare e.g. CD79b, ICOSLG, CD86, CD21, IL-4R in Figure 2D). In contrast, we observed mild phenotypical differences between resting adult and cord blood subsets, including significant differential surface expression of PTPRJ, CR2/CD21, CD79b, FCGR2A, IFNGR1 and FCER2 (Figures 3A and S3). Typical surface expression patterns of murine B-1 cells (64) (CD11b/CD18, CD43 or low level of IgD, CD21 or CD23) were not detectable (Figures 3 and S3).

**Figure 3:**
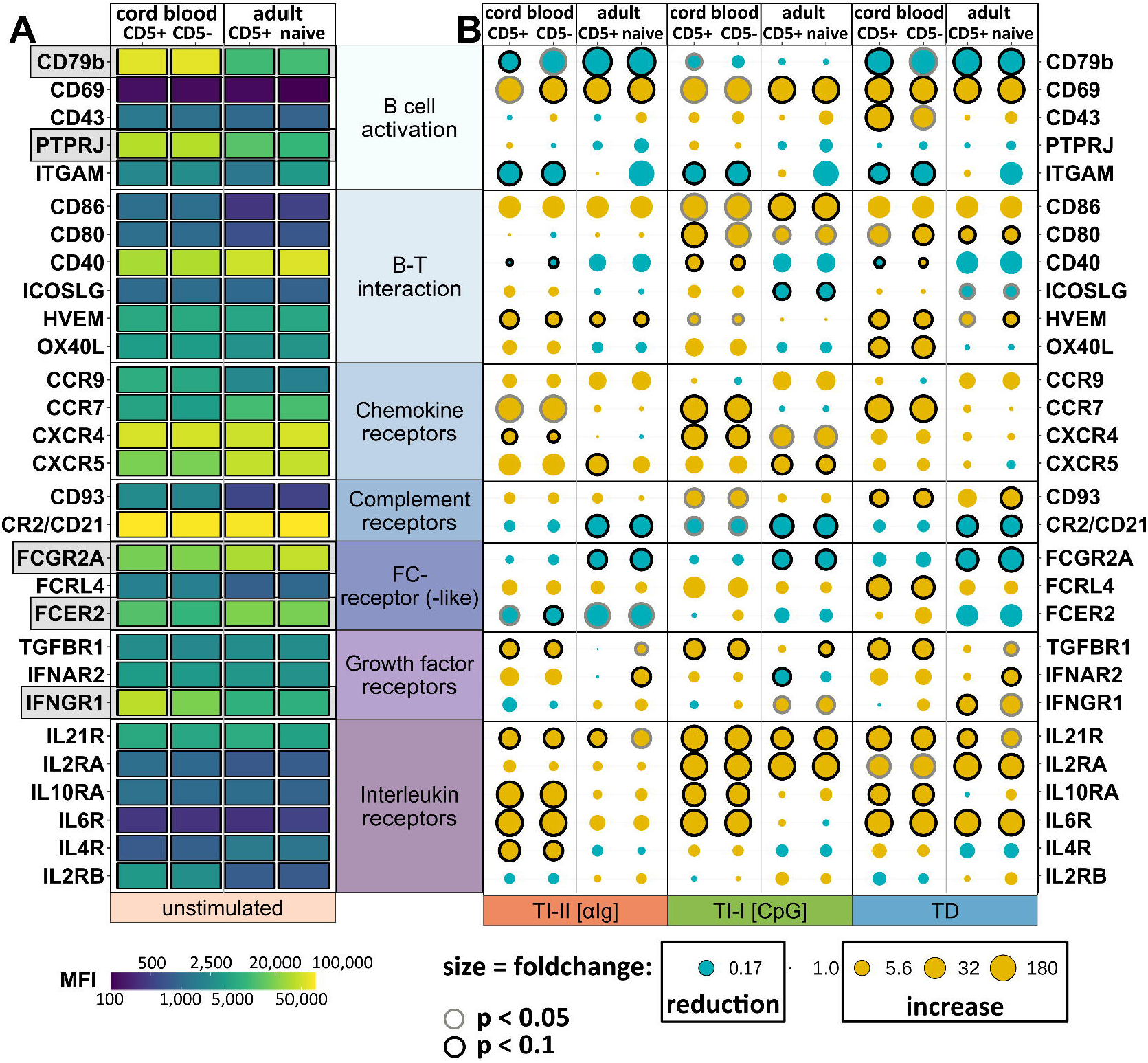
Flow Cytometric Characterization of Cord Blood and Adult Mature B Cells in Resting State and Upon *in vitro* Stimulation. This meta-FACS-analysis is a condensed form of > 4000 FACS plots showing the median MFI values of 29 selected surface markers expressed on cord blood and adult CD19-enriched B cells from three to seven samples for each bar or dot. (A) Heatmap of surface markers in the resting state, grey bars mark significant differences between cord blood and adult mature B cells (p < 0.05, Wilcoxon test). (B) MFI fold-changes after 24 h stimulation under TI-I, TI-II or TD conditions. Dot size and colour code reflect the median MFI fold-change versus unstimulated condition (yellow for increase, blue for decrease). Black (p < 0.05, Wilcoxon test) or grey (p < 0.1, Wilcoxon test) circles show the certainty of increase or decrease compared to resting state. For a graphical description how to read this, see supplemental Figure S3.

We next investigated the responsiveness of mature cord blood and adult naïve B cells *in vitro*, stimulated with CpG (TI-I), anti-Ig (TI-II), and anti-Ig plus CD40L (TD). Compared to steady state, we observed often dramatic changes in surface expression upon stimulation, unidirectional (e.g. CD69, CD86, FCGR2A, CD21) as well as differential (e.g. ITGAM, ICOSLG, OX40L, FCER2, IFNGR, IL4R) between cord blood and adult B cells. The type of stimulus often had a strong impact, mostly on response intensity, with a few exceptions (FCER2, IFNAR2), (Figures 3B and S3). Note that in this analysis CD5^+^ and CD5^-^ B cells were not sort-purified, thus CD5 up- or downregulation during culture cannot be estimated. However, CD5^+^ and CD5^-^ B cells usually shifted homogenously. We conclude that with few exceptions, cord blood and adult mature B cells are phenotypically similar in the resting state but have different response potentials.

### Human Cord Blood B Cell Responses are Rapid but Transient

As human cord blood B cells quickly adopt an activated phenotype upon several types of stimulation, we analysed cord blood and adult B cell response kinetics in more detail and observed further qualitative and quantitative differences. Cord blood B cells proliferated earlier, as the first cell division was already detectable on day 2 upon TI-I or TD stimulation, whereas adult B cells proliferated at the earliest on day 4 (Figure 4A). TI-II stimulation showed weak induction of proliferation in general, in contrast to TD stimulation, particularly in combination with IL-4 and IL-21 (Figures 4B and S4A, respectively). In contrast, adult B cell division was more delayed and barely detectable upon TI stimulation (Figures 4A and B). With regard to the known differences in neonatal and adult T cell composition (2, 4), we performed co-culture assays of mature B cells and autologous T helper cells (CD4^+^CD25^-^), stimulated with anti-CD2/anti-CD3/anti-CD28 beads. The coculture assays confirmed a TD-proliferation potential among both cord blood and adult B cells, although cord blood B cell division was slower than by stimulation with selected ligands (Figure 4B).

**Figure 4:**
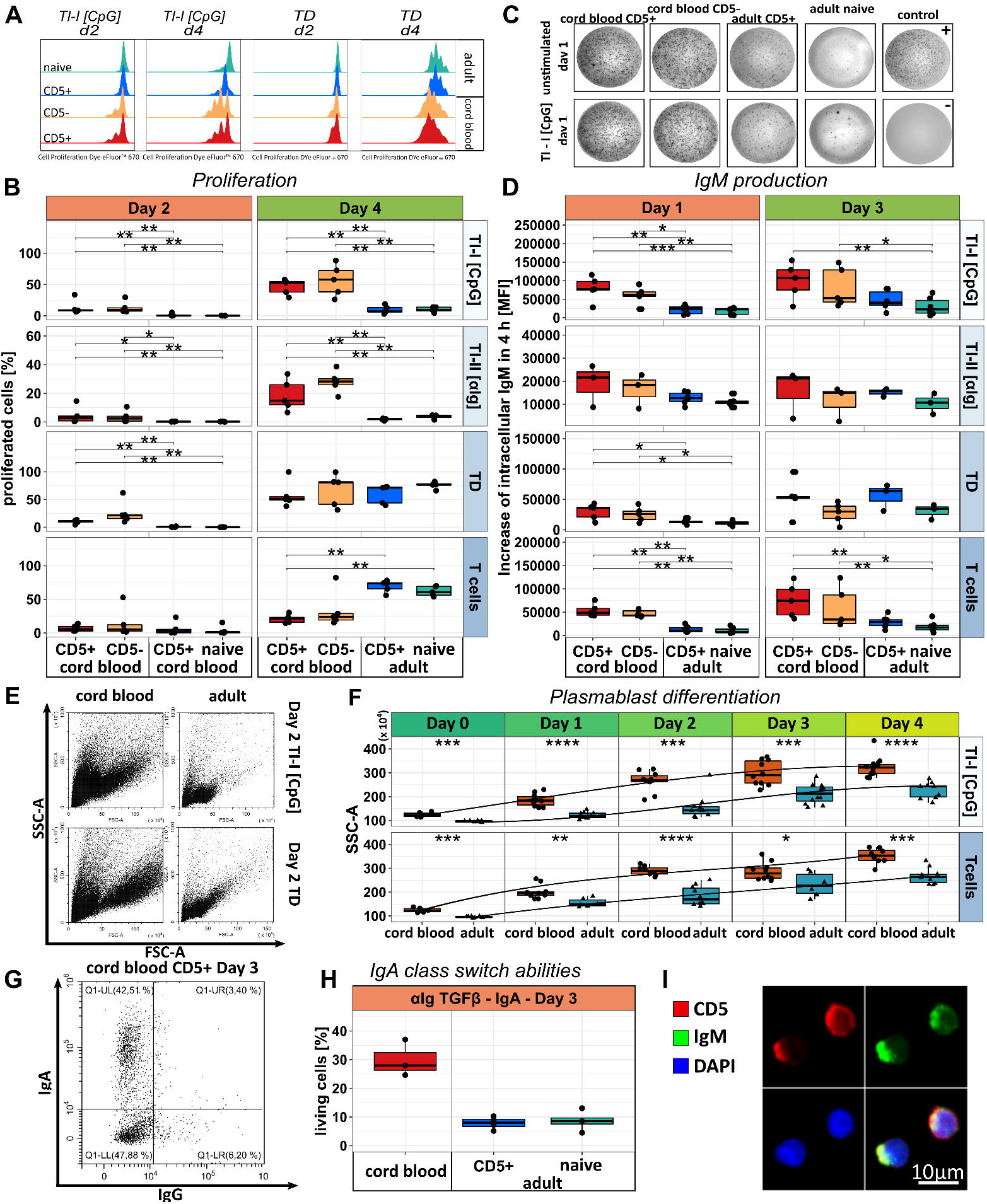
Functional Analysis of Cord Blood and Adult Mature B Cells. (A) Example histograms of the cell proliferation dye eFluor™ 670 in sort-purified cord blood CD5^+^ (red) and CD5^-^ B cells (orange) and adult mature CD5^+^ (blue) and CD5^-^ (mint) B cells until day two (d2) or until day four (d4) upon stimulation with TI-I stimulus [CpG] (left), and TD [anti-Ig + CD40L] (right). (B) Summary of five individual experiments as shown in (A), including TI-II [αIg] stimulation and coculture with activated autologous T cells (CD4^+^CD25^-^). (C) IgM Elispot assays of cord blood and adult mature CD5^+^ and CD5^-^ B cells unstimulated (top) or upon stimulation with TI-I [CpG] (bottom). Plus, and minus indicate positive (IgM memory B cells plus R848) and negative controls (no B cells), respectively. (D) IgM production rate (intracellular IgM accumulation within 4h with GolgiStop™ and GolgiPlug™) of cord blood and adult mature CD5^+^ and CD5^-^ B cells, stimulating conditions as in (B). (E) ASC differentiation as determined by flow cytometry (increasing FSC / SSC) among cord blood and adult B cell samples upon stimulation on day 2. (F) Summary of SSC increase among at least five cord blood or adult B cell samples on d0 to d4 after stimulation (TI-I [CpG], autologous T cells). (G) IgA (ordinate) and IgG (abscissa) class switching of human cord blood B cells on d3 upon stimulation with TGFβ and αIg. (H) Summary of three independent IgA class switching experiments of human cord blood (CD5^+^ and CD5^-^ B cells were not separated for the sake of sufficient cell numbers) and adult CD5^+^ and CD5^-^ B cells, experimental conditions as in (G). (I) Cell polarization (IgM capping) of cord blood mature B cells after 30 minutes of stimulation with CpG. DAPI (blue), CD5 (red), IgM (green). *p < 0.05, **p < 0.01, *** p < 0.001, Wilcoxon rank sum test.

We investigated Ig secretion potentials by ELISpot and showed that cord blood B cells are fully capable of secreting Ig earlier than adult B cells (Figure 4C). Please note, that the ELISpot membrane covered with IgM-capture antibodies likely generates BCR-stimulating signals during the incubation time. Earlier Ig secretion was in line with an increased IgM production rate (Figure 4D), and (in total) accelerated ASC differentiation as determined by FSC/SSC^high^CD38^high^ phenotype (Figures 4E and F and data not shown), PRDM1 induction (Figure S4B), ratio of secreted vs. membrane IgM transcripts (Figure S4C), and cell polarization (“IgM capping”) after 30 minutes of stimulation (Figures 4I and S4F and G) in cord blood vs. adult B cells. In the first four days cord blood B cells homogenously showed ASC differentiation fate, independent of *in vitro* stimulation conditions (Figures 4C, E and F), whereas only a fraction of adult B cells differentiated. This result was consistent with autologous T cell co-culture (Figures 4B, D and F). Neither *in vitro* stimulation nor T cell co-culture induced preferential survival among cord blood over adult B cells, but rather increased apoptosis over time (Figure S4E), supporting the transient nature of cord blood B cell responses.

Finally, as suggested by efficient upregulation of TGFBR1 (Figure 3B), we observed enhanced IgA class switching among cord blood, but not adult B cells treated with anti-Ig and TGFβ, already at day 3 (Figures 4G and H and S4D). In contrast, anti-Ig, CD40L and IL-4 mediated class switching to IgG was more efficient among adult naïve B cells yet delayed until day 5 (Figure S4D).

Our data suggest that cord blood B cells are not impaired, but show rapid TI and TD responsiveness, whereas adult B cell proliferation is mostly confined to TD stimulation. Second, ASC differentiation is the predominant fate of cord blood B cells. These unique dynamics favour transient responses, which may cause the observed quickly waning Ig titres and hyperresponsiveness *in vivo*. The quick and preferential response to TI stimulation, efficient IgM secretion and enhanced IgA class switching are also central to murine B-1 cells (47, 64, 65).

### The Human cord blood BCR Repertoire is Broad, but Inter-individually Conserved

Human cord blood B cells respond fast and efficient to various types of stimulation, however their responses may be hampered by reduced Ig repertoire diversity or skewed IGHV gene usage.

Thus, we performed BCR repertoire deep sequencing of mature CD5^+^ and CD5^-^ B cells from four cord blood and three adult PB samples. We sequenced rearranged IGHV genes of at least 100,000 B cells per population, split into two biological replicates, and used unique molecular identifiers to distinguish clonal expansions from Ig sequence amplification during library generation. The vast majority of BCR rearrangements were unique and unmutated, demonstrating that the cord blood and adult naïve BCR repertoires are similarly variable (Figure 5A). However, whereas clonal expansions were largely absent in adults, they were more frequent in cord blood samples, with almost 2% repeatedly identified sequences on average, and up to 7.3% in one sample (Figures 5A and B). The clonal expansions among both cord blood and adult B cells were devoid of Ig mutations, indicating GC-independent generation. The cord blood-derived clones were larger, up to 40 members, with an average of 4 members per clone (Figure 5C). The comparison of CD5^+^ and CD5^-^ B cell populations within an individual cord blood or adult sample revealed that both subsets were highly related. In any given sample, the majority of clonal expansions were distributed between both subsets, and whereas clones derived from CD5^+^ B cells alone were frequent – particularly in cord blood – clones that consisted only of CD5^-^ B cell-derived sequences were barely detectable (Figures S5A and B), furthermore, indicating that CD5 is not stably expressed on a distinct subset.

**Figure 5:**
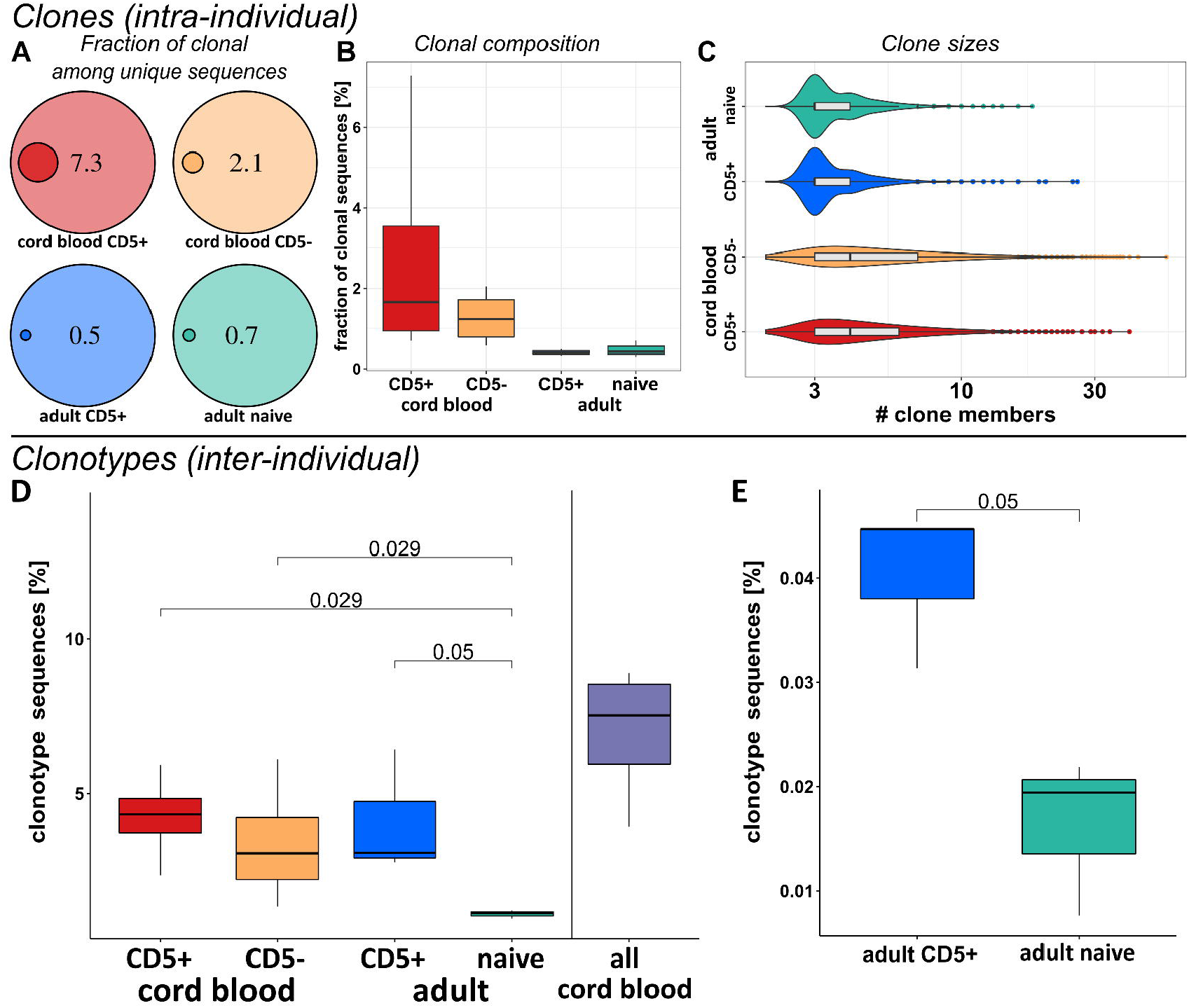
NGS Analysis of the Cord Blood and Adult Mature B Cell Ig Repertoire. Top: Sequences that were repeatedly identified in biological replicates or among different cell samples from the same donor were classified as (int*ra*-individual) clones. (A) Fraction of clonal sequences within one selected sample, and (B) all samples (four cord blood and three adults). (C) Clone sizes detected in all samples. The median clone size among adults is 3 with a maximum of 26 members, whereas the median for cord blood clones is 4 with clone sizes up to 40 members. Bottom: Clonotypes (int*er*-individual): Sequences were classified as clonotypic when bearing the same IGHV gene and at least 90% CDRIII amino acid sequence identity among different donors. (D) The percentage of clonotypes among total cord blood and adult mature CD5^+^ and CD5^-^ B cell derived sequences, and among all cord blood sequences. (E) Number of clonotypes identified in cord blood samples among adult B cell subsets at 100% total amino acid sequence identity. Wilcoxon test was used to determine p-values (four neonates and three adults).

We sought to compare the cord blood BCR repertoire on an inter-individual level and therefore, analysed the distribution of clonotypes, i.e. BCR rearrangements with identical *IGHV* gene and 80% to 100% identical amino acid CDRIII sequence between individuals (Figure S5C). At 90% or more CDRIII identity, cord blood samples (n=4) included 3 to 5% (Ig unmutated) clonotypes and adult naïve B cells (n=3) showed none. In total, 4 to 8% of all BCR rearrangements from four neonates were clonotypic (Figure 5D), suggesting that neonatal B lymphopoiesis includes the selection of conserved specificities. We also observed that adult mature B cells – albeit way less frequent – can show UCB-like BCR specificities, and these clonotypes are significantly enriched among the mature CD5^+^ subset (Figure 5E). Clones and clonotypes barely intersected.

Finally, with regard to murine B-1 cells, the Ig repertoire of human cord blood B cells did not show an enrichment of proximal vs. distal IgV genes, or major biases in IgV gene usage. However, we confirmed a significantly reduced amount of N-nucleotide insertions at D_H_-J_H_-junctions among cord blood B cells (Figure S5D), in line with previous publications (66).

We conclude that the human cord blood mature B cell repertoire is selected to a large extent by inherent mechanisms, which may be similar, but not identical to rodents.

### Human CD34+ hematopoietic precursors from cord blood and adult PB differ in their mature B cell reconstitution capacity in humanized mice

The functional differences between mature B cell subsets from cord blood and adult PB raise the question whether these are simply instructed by e.g. microenvironmental stimulation or cord blood B cells derive from a developmentally distinct lineage, as in mice. We thus, analysed the expression of LET-7b, LIN28b and ARID3A by qRT-PCR among lymphoid precursor cells (CD34^+^) isolated from cord blood and adult PB and observed significant differences, particularly the absence of LIN28b expression in adult precursors (Figure 6A). To determine whether these transcription patterns correlate with their lymphopoietic potential, we generated humanized mice by injecting cord blood (n=16) or adult PB-derived hematopoietic precursor cells (n=8) into NOD.Cg-Prkdc^scid^ Il2rg^tm1Wjl^/SzJ (NSG) mice. Cord blood precursors showed a higher engraftment capability compared to adult-derived cells in the bone marrow microenvironment (Figure 6B). However, the composition of the B cell compartment differed, as cord blood precursors gave rise to a higher frequency of mature CD5+ versus CD5-B cells (Figure 6C), as it is also observed in human cord blood and infant PB (Figure 1D and S1D). Moreover, when normalizing for the variable engraftment efficiency, all mice showed comparable reconstitution of mature B cells in the spleen, but only mice injected with cord blood precursors showed mature (CD5+) B cells in the peritoneal cavity (Figure 6D).

**Figure 6:**
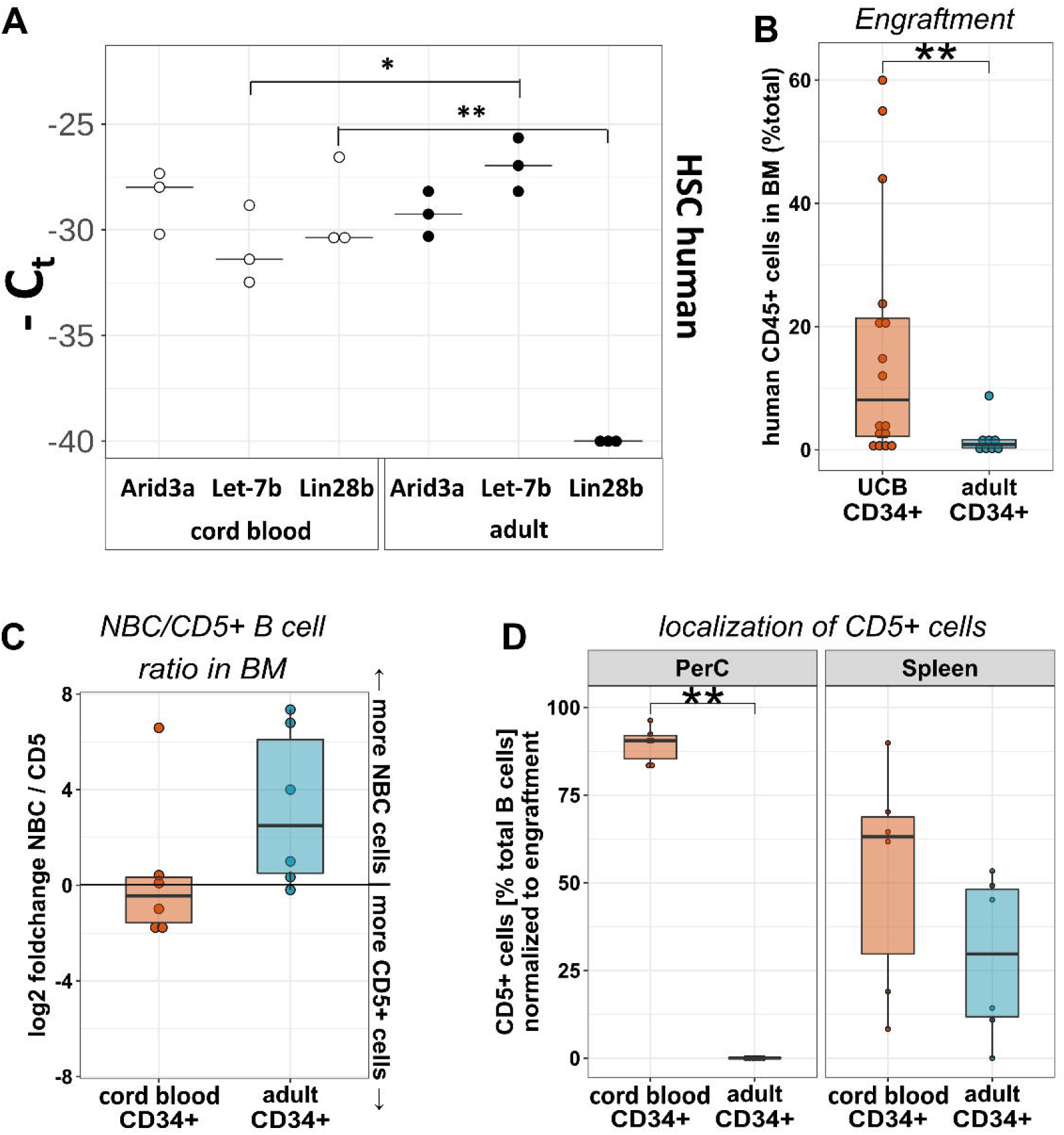
Mature CD5^+^ B Cell Development in Humanized Mice. Flow cytometric analysis of human lymphocytes, derived from NSG mice reconstituted with hematopoietic CD34^+^ progenitor cells from human cord blood or adult BM. (A) Gene expression analysis of selected genes in hematopoietic CD34^+^ progenitor cells from human UCB or adult BM (n>3). *p < 0.05, **p < 0.01, two-sided Wilcoxon test. (B) Engraftment of NSG mice (CD45^+^/total BM mononuclear cells). (C) Ratio of mature CD5^-^ (naïve) versus mature CD5^+^ B cells in the BM of reconstituted mice. (D) Localization of mature CD5^+^ B cells in PerC and spleen of humanized NSG mice, normalized to engraftment.

Human cord blood and adult PB-derived hematopoietic precursor cells differ in their expression of key molecules regulating murine B-1 lineage development, and they provide different engraftment, B cell composition and localization in humanized NSG mice.

## DISCUSSION

Infants suffer from enhanced susceptibility to infections and limited responsiveness to vaccination. Short-lived and quickly waning titres of neutralizing antibody represent a critical factor, particularly during the first months of life. Multiple reasons for this have been described, including immaturity of immune cells or lymphoid tissues. In contrast to previous reports (1–5), our analysis revealed the presence of a large and mature B cell population in cord blood, which responds fast and efficiently *in vitro*. We assume that this population of mature B cells (up to 60% of total cord blood B cells) participates in neonatal and infant humoral immune responses. Recently, it was shown that the neonatal immune system changes rapidly in the first weeks after birth (67), which is likely driven by the extensive host-microbe interactions and potentially also affecting the B cell pool composition. Our data support these data, as cord blood B cells show a strong plasma blast differentiation capacity, thereby favouring changes in the composition of the neonatal immune system in response to a novel environment. However, we show here that the number of mature cells is already high at birth and then largely stable throughout life. We assume that the transition from the cord blood to an adult B cell system occurs slowly but progressively during early childhood, which is in line with the observed slow decrease of CD5+ B cells in children and reflected by the increasing responsiveness and memory formation to various vaccination strategies, which have limited efficiency in neonates and infants.

Our study aimed at a comprehensive characterization of human cord blood B cells to reveal putative differences to adult PB naïve B cells by phenotypical, molecular, and functional analysis. Cord blood B cells show a mature phenotype, but express higher levels of IgM and CD38), which may also contribute to their fast responsiveness, even superior to that of memory B cells (57, 68, 69). On transcriptional level, a major distinctness can be observed, spanning thousands of differentially expressed transcripts and gene sets –, suggesting that neonatal B cell responses differ from those of adult B cells, particularly in their responsiveness to TI versus TD stimulation. Indeed, functional analyses revealed that neonatal B cells are highly sensitive to TI and TD stimulation, whereas adult B cells required T cell help for best performances. The reported reduction in class switching (17, 18) is putatively limited by cytokine levels in neonates (70), but not intrinsic to cord blood B cells, as these showed fast and efficient CSR in particular to IgA. We suggest that the high propensity to IgA (over IgG) class switching supports the quick generation of mucosal immunity in neonates, again reflecting that already at birth mature B cells exist in humans that actively – and quickly – help the immune system to adapt to the environmental challenges. The human neonatal BCR repertoire is unmutated, but highly variable and in this regard no major differences to the adult repertoire could be observed. However, we observed clonal expansions lacking intraclonal diversification, indicating that early in life selection processes exist, that do not require GC diversification. Such clonal expansions were also observed among adult B cells, and then always related to the mature CD5^+^ B cell subset, showing autoimmune specificities (71, 72), associated with autoimmune disorders (73, 74), and chronic lymphocytic leukaemia pathogenesis (52). It was previously reported that neonates show few similar Ig rearrangements (clonotypes) (75), however our dataset contains up to 8% of inter-individually conserved BCR rearrangements among cord blood B cells. It is tempting to speculate on selection of autoantigenic specificities during neonatal B lymphopoiesis, similar to mouse models (76) and potentially contributing to removal of cell debris in the periphery of human neonates (47). Presumably, conserved clonotypes in the cord blood B cell repertoire cause the consistent presence of IgM autoantibodies in the serum of neonates (77). The rarity of cord blood clonotypes among adult naïve B cells raises the question whether these rare clones are long-lived or adult-derived, and whether *de-novo* generated BCR specificities instruct a neonatal B cell fate (36). We propose that the neonatal mature B cell repertoire is substantially selected by inherent, evolutionary conserved mechanisms, likely to a large extent including selective autoimmunity as detectable in serum antibody specificities in cord blood (77).

Our study shows that these specifically selected mature B cells are highly sensitive to stimulation and quickly differentiate into ASCs. However, considering the immature infrastructure of lymphoid tissues (22–24), the rarity of PC survival niches in the bone marrow (25, 26), and the deprivation of survival signals (28), it is easily anticipated why the rapid cord blood B cell responses are transient and wane quickly (3, 5). Moreover, in contrast to adult naïve B cells, cord blood B cells themselves repeatedly showed a bias to undergo apoptosis upon stimulation in vitro. Taken together, we assume that the quick responsiveness of cord blood B cells provides a suitable primary defence against a broad number of infectious agents. However, the transient dynamics then limit this primary response potential, as reactive cells are quickly depleted and leave new-born humans hyperresponsive to excess or too quickly repeated infections (2, 3, 5).

The reason why cord blood B cells behave so distinct from their adult counterpart is difficult to answer in studies on humans. It could be that the distinctness is simply imposed, e.g. by a unique microenvironment in neonates and infants. However, it has been shown that human hematopoietic progenitor cells from cord blood exhibit higher differentiation potential and repopulation capacity compared to adult precursors (78), and it was suggested that human cord blood progenitor subsets diverge in B cell lineage development potential from that of adults (79–81). In line with these findings, the reconstitution of NSG mice with cord blood versus adult-derived CD34^+^ precursors revealed differences in in engraftment efficiency and reconstitution potential of the B cell compartment. This shows that under similar microenvironmental conditions, B cell development from cord blood precursors is qualitatively (and quantitatively) different from that of adult precursors. In line with the characteristic expression pattern in murine B-1 cell development, human cord blood, but not adult precursors express LIN28b, and the downstream expression of the inhibitory micro-RNA LET-7b shifted significantly in favour of the B-1 lineage associated transcription factor ARID3A. We suggest that besides the developmental program, also the conserved (autoimmune) selection of the BCR repertoire may be a decisive molecular reason for this. Thus, our study supports the view that the human B cell system is layered as in mice, where B cell lineages with intrinsic differences develop in separate pathways.

Although many aspects of our knowledge on infant B cell immunity is based on mouse models, the human B-1 lineage counterpart is a long-standing matter of controversy. We therefore compared our findings on human cord blood B cells to the murine B-1 lineage. We observed similarities in function, e.g. the fast and high responsiveness to TI stimulation and the fast and efficient IgM secretion of cord blood B cells argues for overlapping functions. However, the murine B-1 cell-typical transcriptional patterns were barely detectable in humans, and the cord blood BCR repertoire did not show the biased Ig gene usage observed in mice. Thus, human neonatal B cells can only partially be explained by murine B-1 lineage characteristics.

We conclude that human cord blood and adult naive B cells are phenotypically similar but differ markedly in their response potentials. This major distinctness exists in mice and humans, suggesting that a fast and independent IgM and IgA antibody response early in life proved beneficial in evolution. However, the molecular patterns underlying these neonatal B cell responses vary between species. Our study contrasts with many aspects of the current view on human neonatal and infant B cell responses. This has important implications not only for our understanding of infant vulnerability to microbes, but also may explain why certain vaccination strategies rather cause hyperresponsiveness than long-lasting protection. Consequently, systematic vaccination strategies encompassing the neonate’s environment (elder siblings and insufficiently immunized adults) are mandatory to generate herd immunity. Moreover, passive immunization and treatment strategies to sustain ASC survival should be taken into research focus.

## Materials and Methods

### Human samples

PB samples from healthy adults and cord blood samples from healthy term neonates (n > 200, ratio male / female 0.97) were obtained after informed consent according to the Declaration of Helsinki, and approval by the ethics committee of the Medical Faculty at the University of Duisburg-Essen, Germany (BO-10-4380). For samples from PB of children aged 1.5 to 6 years, the study was approved by the Human Research Ethics Committee of the Medical Faculty at University of Gothenburg, Sweden. Informed written consent was obtained from the parents of the children.

### Humanized Mice

NOD.Cg-Prkdc^scid^ Il2rg^tm1Wjl^/SzJ (NSG) mice (https://www.jax.org/strain/005557), were bred and housed at the University Hospital Essen animal care facility. All animal experiments were carried out following institutional guidelines and with protocols approved by the Animal Care Committee of the University Hospital Essen. All mice used in the experiments were kept single housed under pathogen-free conditions. In total 24 NSG mice (12 male and 12 female, age 8-14 weeks, weight 31 ± 5 g) were treated intravenously with 30 mg/kg Busulfan (Busilvex, Pierre Fabre, Sankt Georgen) 24 hours prior to cell transfer, were transplanted by injection into tail-vein with approx. 200,000 human CD34+ progenitor cells derived from UCB or adult PB. Between 2 and 3 months after injection, blood serum, spleen and BM were harvested and a PerC lavage was performed. Harvested tissues were homogenized, splenic erythrocytes were lysed and mononuclear cells extracted by density gradient centrifugation (Ficoll).

### Magnetic cell separation

Human lymphocytes or hematopoietic precursor cells were isolated by Ficoll density centrifugation (density 1.077 g/ml, Pan BioTech, Aidenbach, Germany) followed by selection of CD19-, CD3- or CD34-expressing cells by magnetic cell separation (MACS) (Miltenyi Biotec, Bergisch Gladbach, Germany).

### Flow cytometry

Lymphocytes were analysed on a Cytoflex flow cytometer (Beckman Coulter, Krefeld, Germany) using the CytExpert or sort-purified on a FACSAria III cell sorter (BD Biosciences, Heidelberg, Germany) / FACSAria Fusion cell sorter (BD Bioscience) equipped with BD FACSDiva software (BD Biosciences) after staining with fluorescence-conjugated antibodies. If not indicated otherwise, mature CD5^+^ B cells were defined by the marker constellation CD5^+^CD23^+^CD27^-^CD38^low^IgD^+^ and CD5^-^ (naïve) B cells as CD5^-^CD23^+^CD27^-^CD38^low^IgD^+^ cells. Autologous T cells were isolated from CD19-MACS flow through by sort-purification of CD4^+^CD25^-^ lymphocytes. Intracellular IgM-staining (IgM-FITC, BD Biosciences) was performed after the indicated incubation periods using the Fix/Perm-Kit (BD Biosciences) in the presence of Golgi-Stop and Golgi-Plug (both BD Biosciences). *BHLHE41* expression in human B cell subsets was assessed by PrimeFlow RNA assay (Thermo Fisher, Oberhausen, Germany) with high sensitivity Alexa Fluor 647-probe targeting human *BHLHE41* normalized on *CD8A*-probe.

### ELISpot assays

ELISpot membranes (MabTech, Macleod, Australia) were coated with anti-IgM antibodies. B cells were sort-purified and distributed at 5,000 cells in 100 μl RPMI1640 (Pan BioTech) per well. After 16 hours of incubation (37°C and 5% CO_2_), cells were discarded and secreted IgM was visualized (MabTech, Macleod). The developed spots were quantified by an ELISpot reader (ELISpot Reader System, Autoimmun Diagnostika GmbH, Strassberg, Germany).

### In vitro functional assays

Mature B cell subsets were cultured in RPMI 1640 medium with 20% foetal bovine serum (Pan Biotech), 100 U/ml penicillin and 100 μg/ml streptomycin at 37°C and 5% CO_2_. TD stimulation was performed using 0.03 μg/μl anti-immunoglobulin (anti-Ig, Jackson ImmunoResearch, Newmarket, UK) and 1 μg/ml CD40-ligand-HA with 5 ng/ml anti-HA antibodies (R&D Systems, Minneapolis, USA), and separately in combination with IL-4 and IL-21 (100 IU each). T cell independent type 1 (TI-I) stimulation was mimicked by incubation with CpG OND Type B or R848 (InvivoGen, San Diego, USA). TI type 2 (TI-II) simulation was performed by 0.03 μg/μl anti-Ig treatment. For coculture experiments, B cells and autologous T cells were co-cultivated in equal numbers (250,000 cells). T cells were left unstimulated or activated via anti-CD2/anti-CD3/anti-CD28 beads (Treg Suppression Inspector, Miltenyi).

### Survival and proliferation

Survival was determined by assessing the fraction of vital cells at defined time points during culture by flow cytometric exclusion of 4’,6-Diamidin-2-phenylindol (DAPI) (Merck, Darmstadt, Germany) and Annexin V-APC (BD Biosciences) positive cells. To determine proliferation, isolated B cells were stained with Cell Proliferation Dye eFluor™ 670 (5 μM, Invitrogen) and measured after 48 and 96 h.

### Confocal microscopy

For each stimulation condition, 1 x 10^6^ CD3^-^CD10^-^ cells (anti-CD3-APC (BD Biosciences) and anti-CD10-FITC (BD Biosciences)) were sort-purified and stimulated with either TI-II or CpG at 37°C and 5% CO_2_ for 30 min, and subsequently stained with anti-CD5-PE-Cy5 (BD Biosciences, Heidelberg). The stained cells were fixed with Fixation/Permeabilization Solution (BD Biosciences), followed by intracellular staining with anti-IgM FITC (BD Biosciences, Heidelberg) and Hoechst 33342 (Thermo Fisher). Confocal microscopy was performed with a Leica SP8 gSTED Super Resolution Microscope at the Imaging Center Essen (IMCES).

### Transcriptome analysis

IgM memory (CD5^-^CD23^-^CD27^+^CD38^low^IgM^+^IgD^low^) and IgA or IgG class switched memory (CD5^-^CD23^-^ CD27^+^CD38^low^IgG^+^ or IgA^+^) B cells from PB of adult donors, and CD5^-^ (CD5^-^CD23^+^CD27^-^CD38^low^IgD^+^) and CD5^+^ (CD5^+^CD23^+^CD27^-^CD38^low^IgD^+^) B cells from UCB and adult PB, were isolated by cell-sorting (purity >99%) from 6 adult donors and 6 UCB. RNA was isolated by the RNeasy Micro Kit (Qiagen, Hilden, Germany). For the generation of RNAseq libraries, the NuGEN Trio RNA-Seq System (NuGEN, Redwood City, California, USA) was used. Samples were split equally and processed in independent sequencing steps to allow for correction of batch effects. Sequencing was performed with paired-end sequencing and two times 100 bp length. Sequences were aligned with HiSAT2 (version 2.1.0. Kim D, Langmead B and Salzberg SL. HISAT: a fast-spliced aligner with low memory requirements. *Nature Methods* 2015) to the human genome hg38 and analysed with DESeq2,(82) and R (www.R-project.org/). Hierarchical tree analysis was performed using Genespring GX 11 (Agilent Technologies, Waldbronn, Germany).

### Real Time Quantitative (qRT-) or semi-quantitative PCR

qRT-PCR was performed using TaqMan probes (Applied Biosystems) targeting PRDM1 in 9×10^4^ cord blood and adult mature B cell subsets per condition after stimulation for 48 h (unstimulated control, TI-I, TI-II, TD). LIN28B, LET-7b, and ARID3A specific Taqman probes (ABI) were analyzed among >10^6^ human hematopoietic precursors. Total RNA was extracted using RNeasy Micro Kit (Qiagen) and cDNA synthesis was performed according to manufacturer’s instructions using the High-Capacity cDNA Reverse Transcription Kit (Applied Biosystems) for PRDM1, LIN28B and ARID3A, and the Taqman MicroRNA Reverse Transcription Kit (Applied Biosystems) for LET-7b. The qRT-PCR conditions consisted of 2 min at 50°C, 10 min at 95°C and 45 cycles of 15 s at 55°C and 1 min at 60°C. Every experiment was performed three times and the samples were tested in triplicate. The average of ΔCT values for the amplicon of interest was normalized to that of GAPDH. IgM constant-specific primer was 5’-GCCCTGCCCAACAGGGTCA-3’. IgM transcript splice variants for membrane expression or secretion were amplified with 5’-AGCAAAGCAGTGTGGGGTAGA-3’ and 5’-ACACACAGAGCGGCCAGC-3’ primers (all IDT, Leuven, Belgium), respectively. After agarose gelelectrophoresis, products were visualized by GelRed (Biotium, Fremont, USA) and product intensity measured by Software *ImageJ*(83) and normalized to ACTIN amplicon products (5’-GACGACATGGAGAAAATCTG-3’ and 5’-ATGATCTGGGTCATCTTCTC-3’, Sigma-Aldrich, St. Louis, Missouri, USA).

### IGHV gene rearrangement analysis by high throughput sequencing

Genomic DNA was extracted from B cell populations sorted in duplicates (Gentra Puregene Core Kit, Qiagen). Rearranged *IGHV* genes were sequenced by massive parallel sequencing. By PCR using IGHV1, 3 and 4 subgroup primers for *IGHV* framework region 2 and a mixture of all *IGHJ* primers (Supplementary Table 2), DNA was amplified to generate deep sequencing libraries on the MiSeq Illumina platform (Illumina, San Diego, California, USA). In the first two PCR cycles the template (rearranged *IGVH* gene) is copied only once, while 12 nucleotides *unique molecular identifier* (UMI) are introduced into the amplicon by the first set of primers. Following removal of those primers, the constructs are amplified by a second set of primers, which in turn tag the construct with sequences essential for hybridization on the flow cell. Sequencing was performed with custom primers and the rapid run program with paired-end sequencing and two times 300 bp length. Reads generated by MiSeq were included only when the average quality was ≥ 25. Ambiguities between forward and reverse reads were replaced by “N”. Sequences with identical UMI were classified as PCR duplicates and reduced to the longest detected sequence. Each N in a given sequence was replaced by the most frequent (majority vote) nucleotide. Only in-frame sequences detected more than once were further processed. Differences in the amount of sequences between mature CD5^+^ and naive B cells were balanced by random drawing from the population with higher amount of sequences. Sequences were considered clonally related when using the same *IGHV* gene and sharing 100% complementarity determining region III nucleotide sequence identity. All statistical and bioinformatical evaluations were performed in R (www.R-project.org/) and based on the international ImMunoGeneTics information system (IMGT) database (www.imgt.org/). Mutation frequencies were calculated based on the number or relative position of nucleotide exchanges in the *IGHV* region of each sequence in comparison with the most similar allelic variant present in the respective donor (determined from unmutated sequences). *IGHV* identification was carried out with BLAST. *IGHD* and *IGHJ* identification was performed by calculating the best letter-wise matching gene. For IGH gene usage, the fraction of a given IGH gene among total sequences (all subgroups) detected was calculated. Every nucleotide between and including the first and the last non-germline encoded nucleotide between *IGHV* and *IGHD* or *IGHD* and *IGHJ* gene, with highest germline identity was counted as N1 (V-D) or N2 (D-J) non-germline encoded nucleotide, respectively. IGHV usage was determined by R-package iggeneusage (84).

### Statistical Analysis

Statistical parameters including the description of each data point (n value), the number of mice or human samples per experiment, the number of replicates, the meaning of bars and the statistical tests used are contained in the figure legends. Statistics were calculated using R (www.R-project.org/). Other R packages or software used are given in the related section. For comparisons between two groups a two-sided paired Wilcoxon-test was used. For comparison between more than two groups statistical analysis was performed using a one-way ANOVA with the post-hoc Tukey test. In the figures, asterisks represent p-values: *p < 0.05, **p < 0.01, ***p < 0.001, ****p < 0.0001.

## Author Contributions

M.S. developed the concept and prepared the manuscript. B.B., A.K. M.B., and E.H. designed and carried out most of the experiments and contributed to manuscript preparation. B.B. developed and performed bioinformatical and statistical analysis. K.B., J. A. R., A.G., M.A.W., J.D., T.K., S.V., F.M., P.J., A-C.L. and A.R. contributed experimental work and data interpretation. P.R., D.A. and P.K. contributed samples. M.L., W.H., J.D., and B.G. designed experiments and contributed valuable discussion. R.K., C.C.O., M.A.W., B.B., D.H., B.G., A-C.L. and A.R. reviewed the manuscript.

## Data availability

The *IGHV* gene sequences have been deposited in the Genbank Sequence Read Archive under accession number SRP142713. RNA Sequencing data is available from the GEO database under accession no. GSEXXXX.

## Acknowledgments

We thank Julia Jesdinsky-Elsenbruch for excellent technical assistance and Klaus Lennartz for his valuable engineering support. We thank Ludger Klein-Hitpass and the staff members at the Biochip-laboratory at the Institute of Cell Biology in Essen. We thank the staff members at the flow cytometry and fluorescence core facility Imaging Center Essen (IMCES). This work was supported by the Deutsche Forschungsgemeinschaft (grants SE1885/2-1, Ku1315/10-1, TRR60, DU1964/1-1), the Deutsche Krebshilfe (grant 70112628), the Swedish Research Council (grant 2017-01118), and Cancerfonden (grant CAN 2018/710).

## References

1. B. Adkins, C. Leclerc, S. Marshall-Clarke, Neonatal adaptive immunity comes of age. Nat Rev Immunol 4, 553–564 (2004).

2. T. R. Kollmann, B. Kampmann, S. K. Mazmanian, A. Marchant, O. Levy, Protecting the Newborn and Young Infant from Infectious Diseases: Lessons from Immune Ontogeny. Immunity 46, 350–363 (2017).

3. C. A. Siegrist, R. Aspinall, B-cell responses to vaccination at the extremes of age. Nat Rev Immunol 9, 185–194 (2009).

4. O. Levy, Innate immunity of the newborn: basic mechanisms and clinical correlates. Nat Rev Immunol 7, 379–390 (2007).

5. A. J. Pollard, K. P. Perrett, P. C. Beverley, Maintaining protection against invasive bacteria with protein-polysaccharide conjugate vaccines. Nat Rev Immunol 9, 213–220 (2009).

6. D. H. Smith, G. Peter, D. L. Ingram, A. L. Harding, P. Anderson, Responses of children immunized with the capsular polysaccharide of Hemophilus influenzae, type b. Pediatrics 52, 637–644 (1973).

7. G. Blanchard-Rohner, A. S. Pulickal, C. M. Jol-van der Zijde, M. D. Snape, A. J. Pollard, Appearance of peripheral blood plasma cells and memory B cells in a primary and secondary immune response in humans. Blood 114, 4998–5002 (2009).

8. D. F. Kelly et al., CRM197-conjugated serogroup C meningococcal capsular polysaccharide, but not the native polysaccharide, induces persistent antigen-specific memory B cells. Blood 108, 2642–2647 (2006).

9. R. Mitchell, D. F. Kelly, A. J. Pollard, J. Truck, Polysaccharide-specific B cell responses to vaccination in humans. Hum Vaccin Immunother 10, 1661–1668 (2014).

10. R. Booy et al., Immunogenicity of combined diphtheria, tetanus, and pertussis vaccine given at 2, 3, and 4 months versus 3, 5, and 9 months of age. Lancet 339, 507–510 (1992).

11. I. Debock, V. Flamand, Unbalanced Neonatal CD4(+) T-Cell Immunity. Front Immunol 5, 393 (2014).

12. A. W. Griffioen, S. W. Franklin, B. J. Zegers, G. T. Rijkers, Expression and functional characteristics of the complement receptor type 2 on adult and neonatal B lymphocytes. Clin Immunol Immunopathol 69, 1–8 (1993).

13. K. Kaur, S. Chowdhury, N. S. Greenspan, J. R. Schreiber, Decreased expression of tumor necrosis factor family receptors involved in humoral immune responses in preterm neonates. Blood 110, 2948–2954 (2007).

14. M. A. Pettengill, O. Levy, Circulating Human Neonatal Naive B Cells are Deficient in CD73 Impairing Purine Salvage. Front Immunol 7, 121 (2016).

15. D. Viemann, P. Schlenke, H. J. Hammers, H. Kirchner, A. Kruse, Differential expression of the B cell-restricted molecule CD22 on neonatal B lymphocytes depending upon antigen stimulation. Eur J Immunol 30, 550–559 (2000).

16. L. Tasker, S. Marshall-Clarke, Functional responses of human neonatal B lymphocytes to antigen receptor cross-linking and CpG DNA. Clin Exp Immunol 134, 409–419 (2003).

17. M. A. Pettengill et al., Distinct TLR-mediated cytokine production and immunoglobulin secretion in human newborn naive B cells. Innate Immun 22, 433–443 (2016).

18. S. Glaesener et al., Decreased production of class-switched antibodies in neonatal B cells is associated with increased expression of miR-181b. PLoS One 13, e0192230 (2018).

19. S. Kruetzmann et al., Human immunoglobulin M memory B cells controlling Streptococcus pneumoniae infections are generated in the spleen. J Exp Med 197, 939–945 (2003).

20. S. Weller et al., Somatic diversification in the absence of antigen-driven responses is the hallmark of the IgM+ IgD+ CD27+ B cell repertoire in infants. J Exp Med 205, 1331–1342 (2008).

21. K. Bauer et al., Homology-directed recombination in IgH variable region genes from human neonates, infants and adults: implications for junctional diversity. Mol Immunol 44, 2969–2977 (2007).

22. M. Pihlgren et al., Unresponsiveness to lymphoid-mediated signals at the neonatal follicular dendritic cell precursor level contributes to delayed germinal center induction and limitations of neonatal antibody responses to T-dependent antigens. J Immunol 170, 2824–2832 (2003).

23. G. Blanchard Rohner et al., The magnitude of the antibody and memory B cell responses during priming with a protein-polysaccharide conjugate vaccine in human infants is associated with the persistence of antibody and the intensity of booster response. J Immunol 180, 2165–2173 (2008).

24. C. Kruschinski, M. Zidan, A. S. Debertin, S. von Horsten, R. Pabst, Age-dependent development of the splenic marginal zone in human infants is associated with different causes of death. Hum Pathol 35, 113–121 (2004).

25. E. Belnoue et al., APRIL is critical for plasmablast survival in the bone marrow and poorly expressed by early-life bone marrow stromal cells. Blood 111, 2755–2764 (2008).

26. A. Marchant et al., Predominant influence of environmental determinants on the persistence and avidity maturation of antibody responses to vaccines in infants. J Infect Dis 193, 1598–1605 (2006).

27. M. Pihlgren et al., Delayed and deficient establishment of the long-term bone marrow plasma cell pool during early life. Eur J Immunol 31, 939–946 (2001).

28. M. Pihlgren et al., Reduced ability of neonatal and early-life bone marrow stromal cells to support plasmablast survival. J Immunol 176, 165–172 (2006).

29. K. G. Smith, A. Light, G. J. Nossal, D. M. Tarlinton, The extent of affinity maturation differs between the memory and antibody-forming cell compartments in the primary immune response. EMBO J 16, 2996–3006 (1997).

30. K. Hayakawa, R. R. Hardy, L. A. Herzenberg, L. A. Herzenberg, Progenitors for Ly-1 B cells are distinct from progenitors for other B cells. J Exp Med 161, 1554–1568 (1985).

31. A. M. Hamilton, J. F. Kearney, Effects of IgM allotype suppression on serum IgM levels, B-1 and B-2 cells, and antibody responses in allotype heterozygous F1 mice. Dev Immunol 4, 27–41 (1994).

32. N. A. Mabbott, D. Gray, Identification of co-expressed gene signatures in mouse B1, marginal zone and B2 B-cell populations. Immunology 141, 79–95 (2014).

33. B. Vilagos et al., Essential role of EBF1 in the generation and function of distinct mature B cell types. J Exp Med 209, 775–792 (2012).

34. Z. Xu et al., Cyclin-dependent kinase inhibitor Cdkn2c regulates B cell homeostasis and function in the NZM2410-derived murine lupus susceptibility locus Sle2c1. J Immunol 186, 6673–6682 (2011).

35. K. P. Lam, K. Rajewsky, B cell antigen receptor specificity and surface density together determine B-1 versus B-2 cell development. J Exp Med 190, 471–477 (1999).

36. R. Graf et al., BCR-dependent lineage plasticity in mature B cells. Science 363, 748–753 (2019).

37. R. M. Perlmutter, J. F. Kearney, S. P. Chang, L. E. Hood, Developmentally controlled expression of immunoglobulin VH genes. Science 227, 1597–1601 (1985).

38. L. Carlsson, D. Holmberg, Genetic basis of the neonatal antibody repertoire: germline V-gene expression and limited N-region diversity. Int Immunol 2, 639–643 (1990).

39. A. B. Kantor, C. E. Merrill, L. A. Herzenberg, J. L. Hillson, An unbiased analysis of V(H)-D-J(H) sequences from B-1a, B-1b, and conventional B cells. J Immunol 158, 1175–1186 (1997).

40. Y. Yang et al., Distinct mechanisms define murine B cell lineage immunoglobulin heavy chain (IgH) repertoires. Elife 4, e09083 (2015).

41. K. Hayakawa, R. R. Hardy, D. R. Parks, L. A. Herzenberg, The “Ly-1 B” cell subpopulation in normal immunodefective, and autoimmune mice. J Exp Med 157, 202–218 (1983).

42. F. Caligaris-Cappio, M. Gobbi, M. Bofill, G. Janossy, Infrequent normal B lymphocytes express features of B-chronic lymphocytic leukemia. J Exp Med 155, 623–628 (1982).

43. H. Wardemann, T. Boehm, N. Dear, R. Carsetti, B-1a B cells that link the innate and adaptive immune responses are lacking in the absence of the spleen. J Exp Med 195, 771–780 (2002).

44. D. O. Griffin, N. E. Holodick, T. L. Rothstein, Human B1 cells in umbilical cord and adult peripheral blood express the novel phenotype CD20+ CD27+ CD43+ CD70. J Exp Med 208, 67–80 (2011).

45. C. A. Reynaud, J. C. Weill, Gene profiling of CD11b(+) and CD11b(-) B1 cell subsets reveals potential cell sorting artifacts. J Exp Med 209, 433–434 (2012).

46. M. Perez-Andres et al., The nature of circulating CD27+CD43+ B cells. J Exp Med 208, 2565–2566 (2011).

47. N. Baumgarth, The double life of a B-1 cell: self-reactivity selects for protective effector functions. Nat Rev Immunol 11, 34–46 (2011).

48. M. Rakhmanov et al., Circulating CD21low B cells in common variable immunodeficiency resemble tissue homing, innate-like B cells. Proc Natl Acad Sci U S A 106, 13451–13456 (2009).

49. G. R. Ehrhardt et al., Expression of the immunoregulatory molecule FcRH4 defines a distinctive tissue-based population of memory B cells. J Exp Med 202, 783–791 (2005).

50. T. Kreslavsky, J. B. Wong, M. Fischer, J. A. Skok, M. Busslinger, Control of B-1a cell development by instructive BCR signaling. Curr Opin Immunol 51, 24–31 (2018).

51. D. J. DiLillo, T. Matsushita, T. F. Tedder, B10 cells and regulatory B cells balance immune responses during inflammation, autoimmunity, and cancer. Ann N Y Acad Sci 1183, 38–57 (2010).

52. M. Seifert et al., Cellular origin and pathophysiology of chronic lymphocytic leukemia. J Exp Med 209, 2183–2198 (2012).

53. G. P. Sims et al., Identification and characterization of circulating human transitional B cells. Blood 105, 4390–4398 (2005).

54. H. H. Wortis, M. Teutsch, M. Higer, J. Zheng, D. C. Parker, B-cell activation by crosslinking of surface IgM or ligation of CD40 involves alternative signal pathways and results in different B-cell phenotypes. Proc Natl Acad Sci U S A 92, 3348–3352 (1995).

55. E. Blanco et al., Age-associated distribution of normal B-cell and plasma cell subsets in peripheral blood. J Allergy Clin Immunol 141, 2208–2219 e2216 (2018).

56. H. Morbach, E. M. Eichhorn, J. G. Liese, H. J. Girschick, Reference values for B cell subpopulations from infancy to adulthood. Clin Exp Immunol 162, 271–279 (2010).

57. M. Seifert et al., Functional capacities of human IgM memory B cells in early inflammatory responses and secondary germinal center reactions. Proc Natl Acad Sci U S A 112, E546–555 (2015).

58. A. Cerutti, The regulation of IgA class switching. Nat Rev Immunol 8, 421–434 (2008).

59. J. Yuan, C. K. Nguyen, X. Liu, C. Kanellopoulou, S. A. Muljo, Lin28b reprograms adult bone marrow hematopoietic progenitors to mediate fetal-like lymphopoiesis. Science 335, 1195–1200 (2012).

60. Y. Zhou et al., Lin28b promotes fetal B lymphopoiesis through the transcription factor Arid3a. J Exp Med 212, 569–580 (2015).

61. T. Kreslavsky et al., Essential role for the transcription factor Bhlhe41 in regulating the development, self-renewal and BCR repertoire of B-1a cells. Nat Immunol 18, 442–455 (2017).

62. F. Edfors et al., Gene-specific correlation of RNA and protein levels in human cells and tissues. Mol Syst Biol 12, 883 (2016).

63. M. Gry et al., Correlations between RNA and protein expression profiles in 23 human cell lines. BMC Genomics 10, 365 (2009).

64. A. B. Kantor, L. A. Herzenberg, Origin of murine B cell lineages. Annu Rev Immunol 11, 501–538 (1993).

65. A. J. Macpherson et al., A primitive T cell-independent mechanism of intestinal mucosal IgA responses to commensal bacteria. Science 288, 2222–2226 (2000).

66. B. Hong et al., In-Depth Analysis of Human Neonatal and Adult IgM Antibody Repertoires. Front Immunol 9, 128 (2018).

67. A. Olin et al., Stereotypic Immune System Development in Newborn Children. Cell 174, 1277–1292 e1214 (2018).

68. N. Engels, J. Wienands, Memory control by the B cell antigen receptor. Immunol Rev 283, 150–160 (2018).

69. Y. Shi, K. Agematsu, H. D. Ochs, K. Sugane, Functional analysis of human memory B-cell subpopulations: IgD+CD27+ B cells are crucial in secondary immune response by producing high affinity IgM. Clin Immunol 108, 128–137 (2003).

70. M. Nesin, S. Cunningham-Rundles, Cytokines and neonates. Am J Perinatol 17, 393–404 (2000).

71. R. R. Hardy, K. Hayakawa, M. Shimizu, K. Yamasaki, T. Kishimoto, Rheumatoid factor secretion from human Leu-1+ B cells. Science 236, 81–83 (1987).

72. M. Herve et al., Unmutated and mutated chronic lymphocytic leukemias derive from self-reactive B cell precursors despite expressing different antibody reactivity. J Clin Invest 115, 1636–1643 (2005).

73. S. E. Burastero, P. Casali, R. L. Wilder, A. L. Notkins, Monoreactive high affinity and polyreactive low affinity rheumatoid factors are produced by CD5+ B cells from patients with rheumatoid arthritis. The Journal of experimental medicine 168, 1979–1992 (1988).

74. M. Dauphinee, Z. Tovar, N. Talal, B cells expressing CD5 are increased in Sjogren’s syndrome. Arthritis and rheumatism 31, 642–647 (1988).

75. C. Soto et al., High frequency of shared clonotypes in human B cell receptor repertoires. Nature 566, 398–402 (2019).

76. K. Hayakawa et al., Positive selection of anti-thy-1 autoreactive B-1 cells and natural serum autoantibody production independent from bone marrow B cell development. J Exp Med 197, 87–99 (2003).

77. Y. Merbl, M. Zucker-Toledano, F. J. Quintana, I. R. Cohen, Newborn humans manifest autoantibodies to defined self molecules detected by antigen microarray informatics. J Clin Invest 117, 712–718 (2007).

78. N. Graffmann et al., Age-Related Increase of EED Expression in Early Hematopoietic Progenitor Cells is Associated with Global Increase of the Histone Modification H3K27me3. Stem Cells Dev 24, 2018–2031 (2015).

79. Y. Hirose et al., B-cell precursors differentiated from cord blood CD34+ cells are more immature than those derived from granulocyte colony-stimulating factor-mobilized peripheral blood CD34+ cells. Immunology 104, 410–417 (2001).

80. E. Sanz, M. Alvarez-Mon, A. C. Martinez, A. de la Hera, Human cord blood CD34+Pax-5+ B-cell progenitors: single-cell analyses of their gene expression profiles. Blood 101, 3424–3430 (2003).

81. E. Sanz et al., Ordering human CD34+CD10-CD19+ pre/pro-B-cell and CD19-common lymphoid progenitor stages in two pro-B-cell development pathways. Proc Natl Acad Sci U S A 107, 5925–5930 (2010).

82. M. I. Love, W. Huber, S. Anders, Moderated estimation of fold change and dispersion for RNA-seq data with DESeq2. Genome Biol 15, 550 (2014).

83. C. A. Schneider, W. S. Rasband, K. W. Eliceiri, NIH Image to ImageJ: 25 years of image analysis. Nat Methods 9, 671–675 (2012).

84. S. Kitanovski, D. Hoffmann, IgGeneUsage: differential gene usage in immune repertoires. Bioinformatics 36, 3590–3591 (2020).

