## Supplemental Figure 1 for "Human neonatal B cell immunity differs from the adult version by conserved Ig repertoires and rapid, but transient response dynamics"

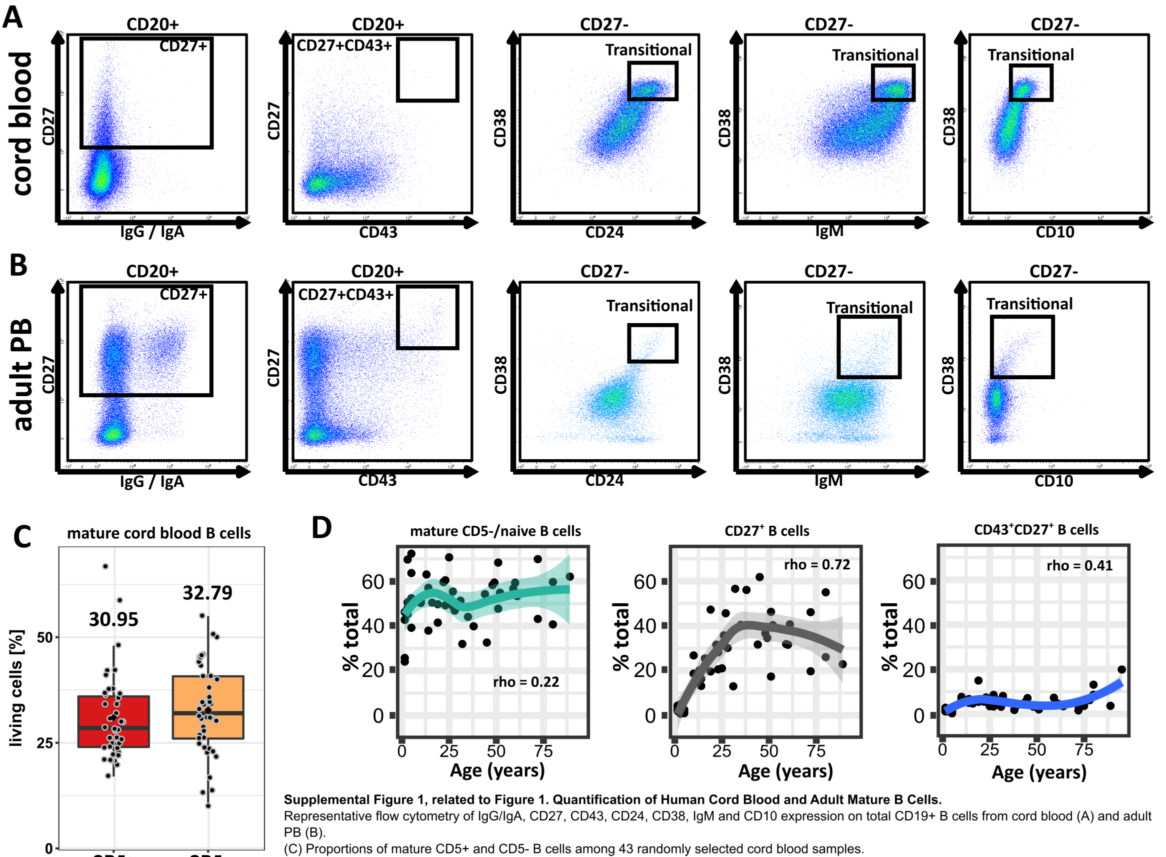


**Supplemental Figure 1, related to Figure 1. Quantification of Human Cord Blood and Adult Mature B Cells.**
Representative flow cytometry of IgG/IgA, CD27, CD43, CD24, CD38, IgM and CD10 expression on total CD19+ B cells from cord blood (A) and adult PB (B).
(C) Proportions of mature CD5+ and CD5- B cells among 43 randomly selected cord blood samples.
(D) Age dependency of mature CD5- (naïve) B cells, CD27+ memory and CD27+CD43+ B cells from 45 donors aged 0 – 90 years. Line of fit (bold line) and 95% confidence interval (shaded area) were calculated by R smooth.
