## Supplemental Figure 2 for "Human neonatal B cell immunity differs from the adult version by conserved Ig repertoires and rapid, but transient response dynamics"

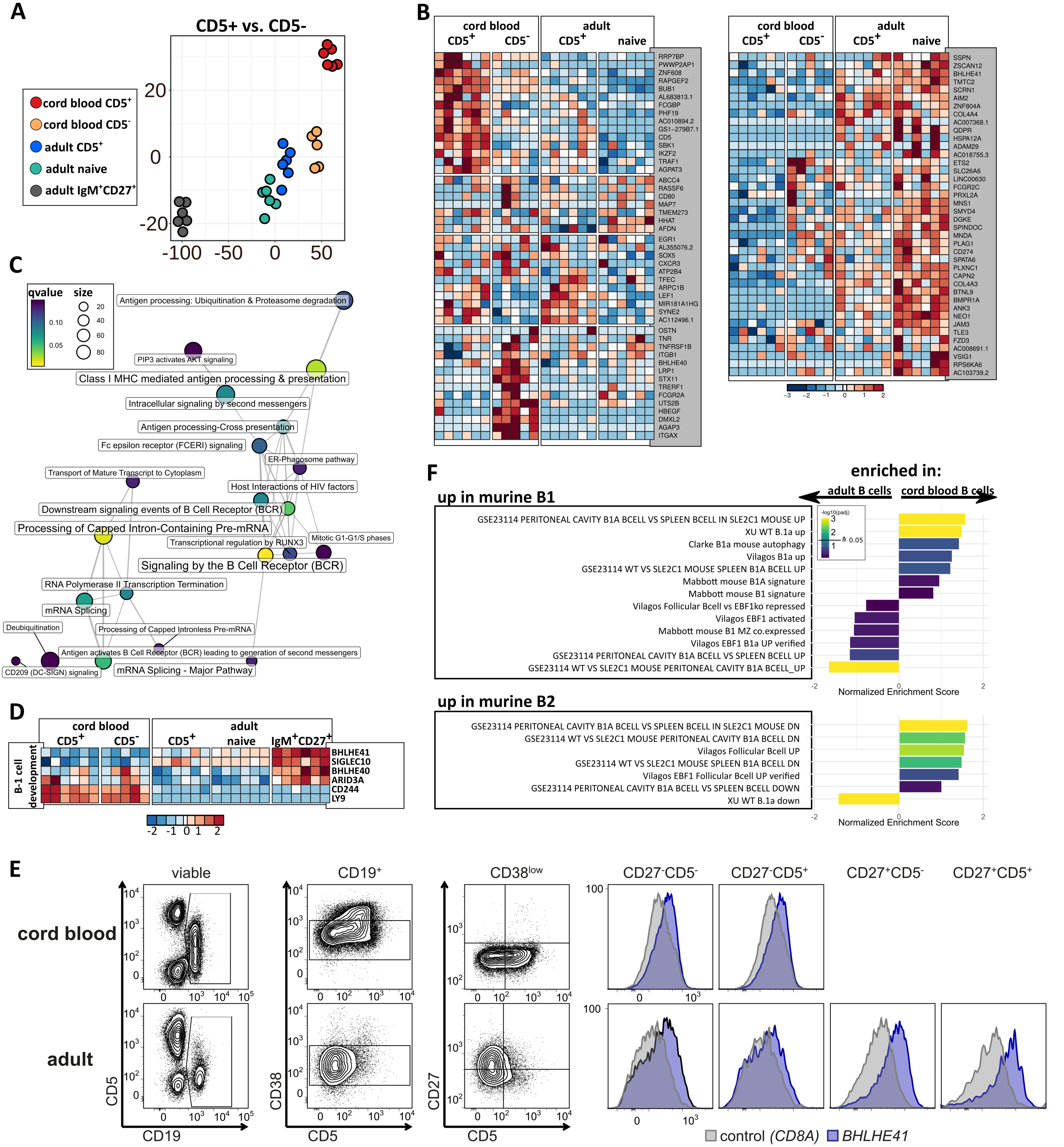


**Supplemental Figure 2, related to Figure 2. Transcriptome Wide Comparison of Cord Blood and Adult B Cells.**

(A) T-SNE plot showing the distribution of cord blood and adult mature CD5+ and CD5- B cell and adult IgM memory B cell subsets, based on 87 significantly different expressed genes (MANOVA, $p < 0.001$, two-fold difference) between CD5+ or CD5- mature B cells, regardless of age status (R-package Rtsne with standard settings).

(B) Detailed heatmap of the 87 genes shown in (A).

(C) Reactome pathway map ("Immune System") of enriched gene sets ($q < 0.15$) between cord blood and adult B cell subsets.

(D) Heatmap for selected genes involved in murine B-1 cell development, expressed among cord blood and adult mature CD5+ and CD5- B cells, and adult IgM memory B cells. The colouring depicts row-wise normalized fold-changes.

(E) BHLHE41 expression in human adult and cord blood B cell subsets as determined by PrimeFlow RNA assay.

(F) GSEA of published murine B-1 or B-2 cell specific gene sets among cord blood versus adult B cells (CD5+ and CD5- pooled).
