## Supplemental Figure 3 for "Human neonatal B cell immunity differs from the adult version by conserved Ig repertoires and rapid, but transient response dynamics"

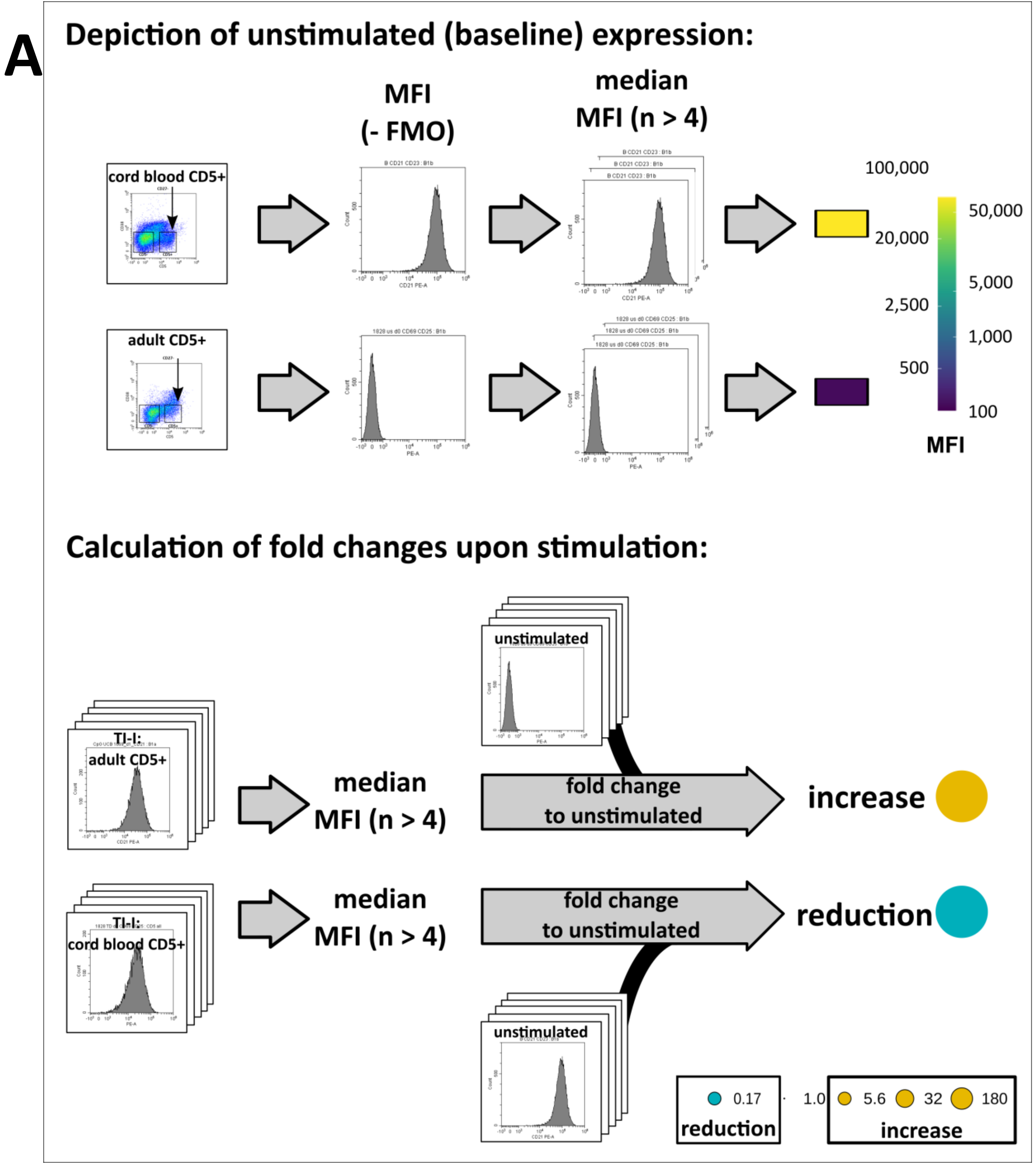


**Supplemental Figure 3. Examples for Flow Cytometry of Cord Blood and Adult Mature B Cells in Resting State and Under *in vitro* Stimulation.** This Figure shows representative flow cytometric data for selected surface molecules underlying the meta-FACS-analysis shown in Figure 3. All flow cytometric data shown in Figures 3 and S3 are based on unstimulated (d0) or stimulated (o/n) samples, that were not sort-purified. Due to the multiple conditions and limited cell numbers per donor, the data shown in each row are not always derived from the same donor. In some instances, contaminating T cells are detectable ($CD5^{high}$), which were excluded by gating strategy. Mean fluorescence intensity (MFI) values are shown in Figure 3, also when in some donors a biphasic distribution was detectable among mature $CD5^{+}$ B cells ($CD5^{dim}$ and $CD5^{high}$) (see e.g. CD79b or IFNGR1). (A) Flow chart of data processing, analysis and representation in Figure 3. (B) Gating strategy for MFI quantification. (C-N) Display of each individual marker included in Figure 3 (Individual plots show Fluorescence minus one control, isotype control, expression at steady state (d0), and TI-I-, and TD-stimulated cells (d1).

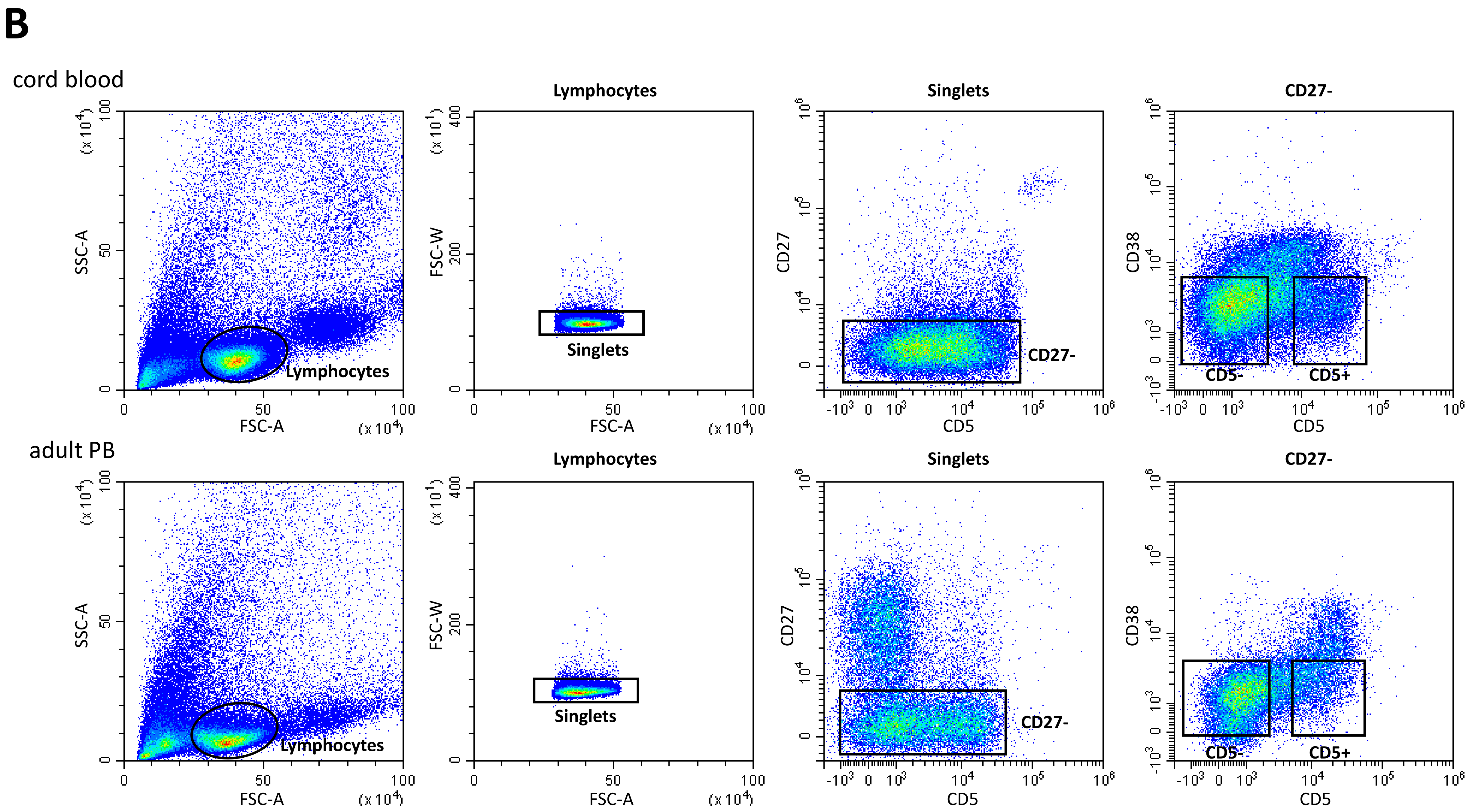
B
cord blood
adult PB
Lymphocytes
Singlets
CD27-
CD5-
CD5+
SSC-A
FSC-A
FSC-W
CD27
CD38
CD5

C

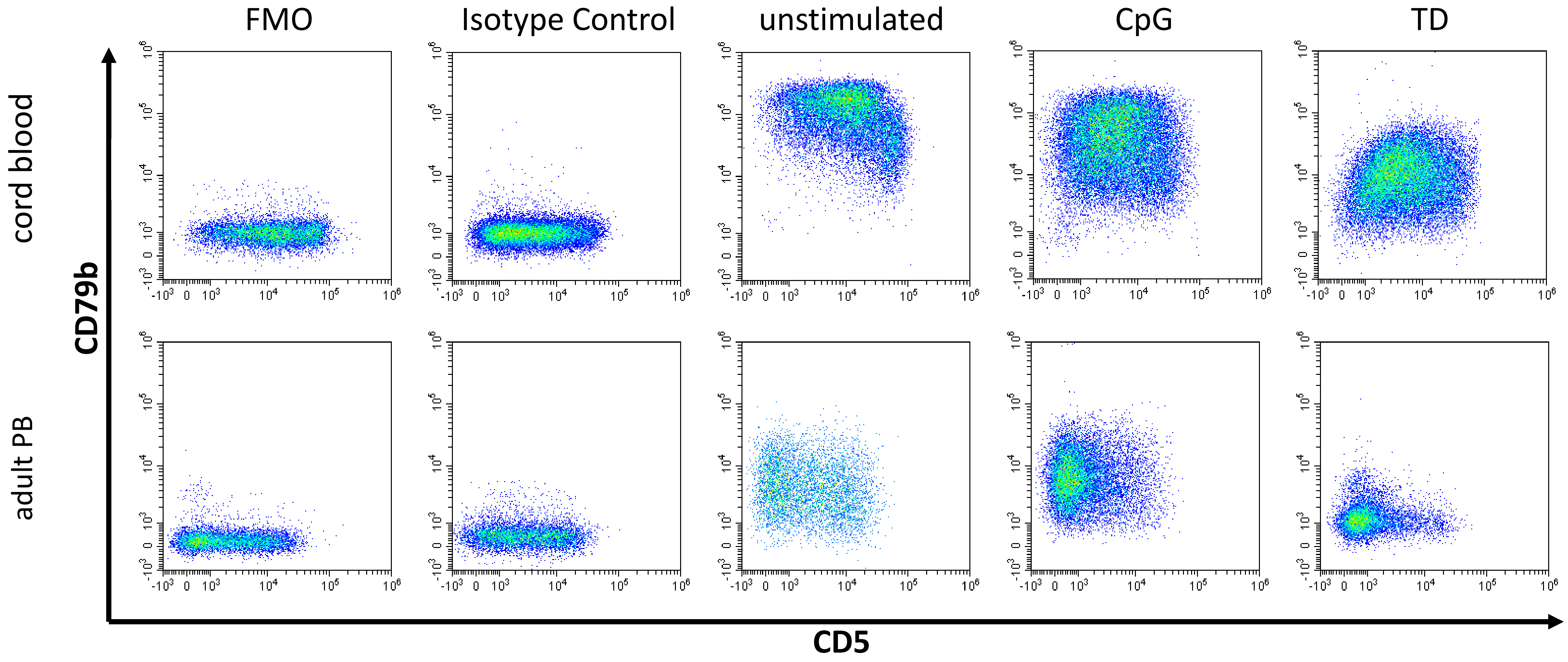
FMO
Isotype Control
unstimulated
CpG
TD
cord blood
adult PB
CD79b
CD5

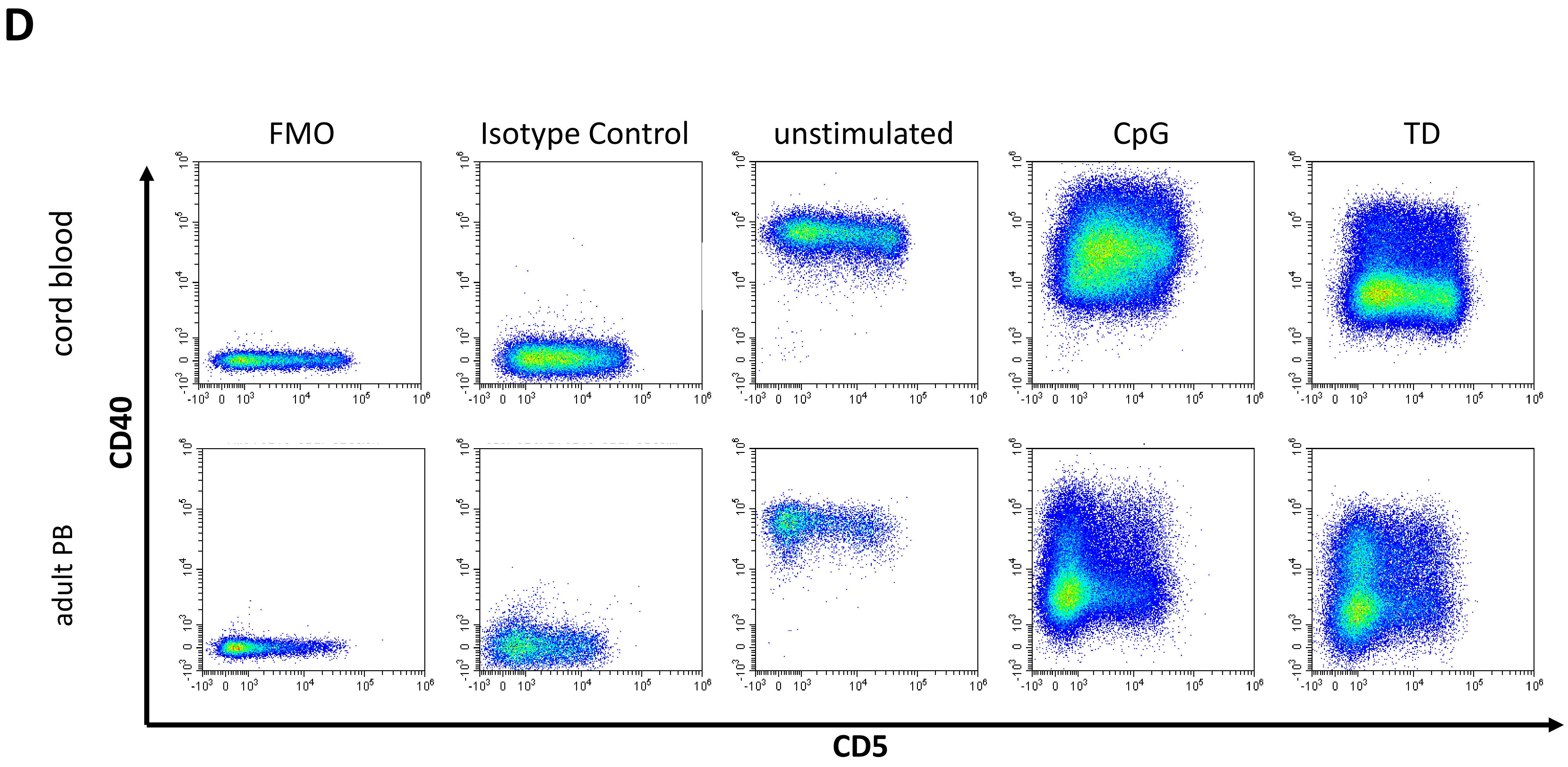
D
FMO
Isotype Control
unstimulated
CpG
TD
cord blood
adult PB
CD40
CD5

E

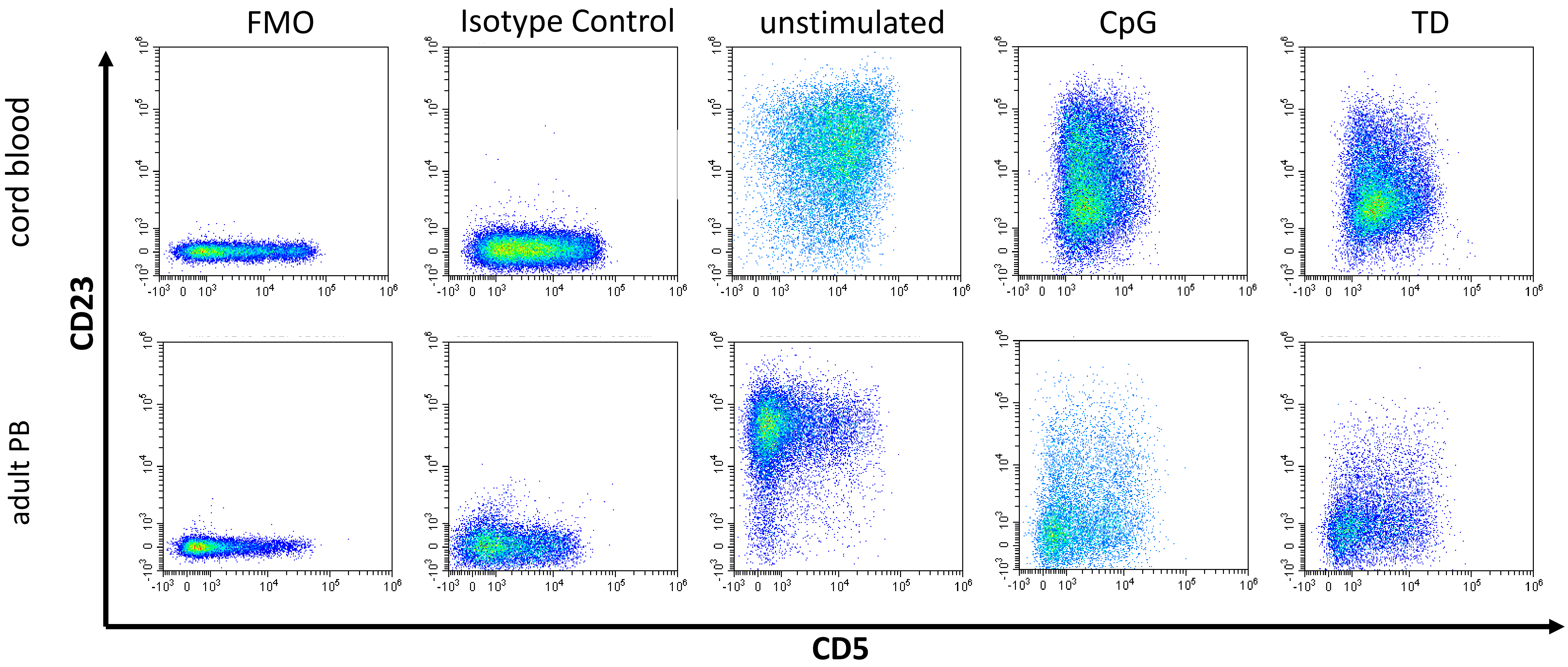

FMO
Isotype Control
unstimulated
CpG
TD
cord blood
adult PB
CD23
CD5

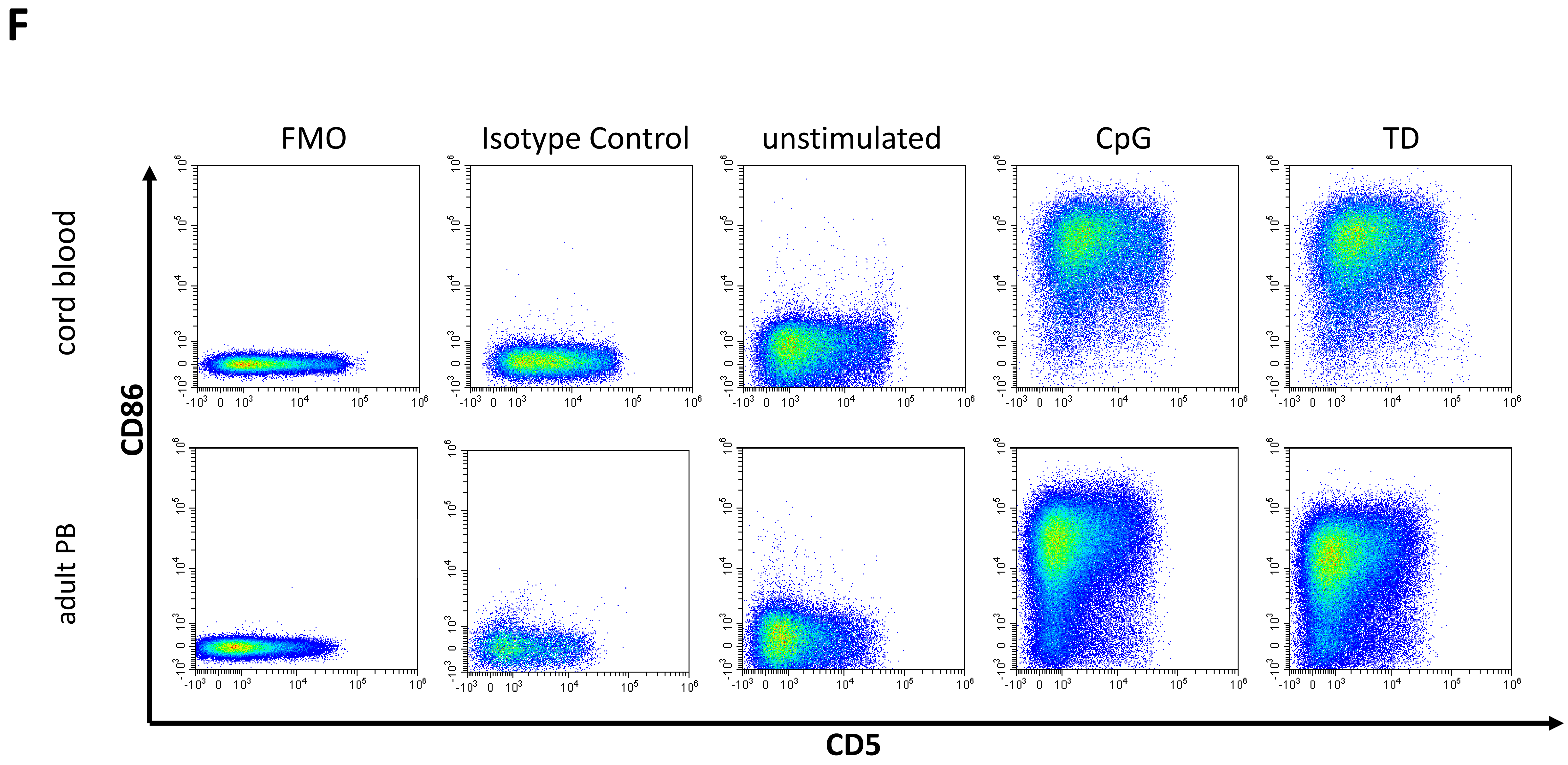
F
FMO
Isotype Control
unstimulated
CpG
TD
cord blood
adult PB
CD86
CD5

G

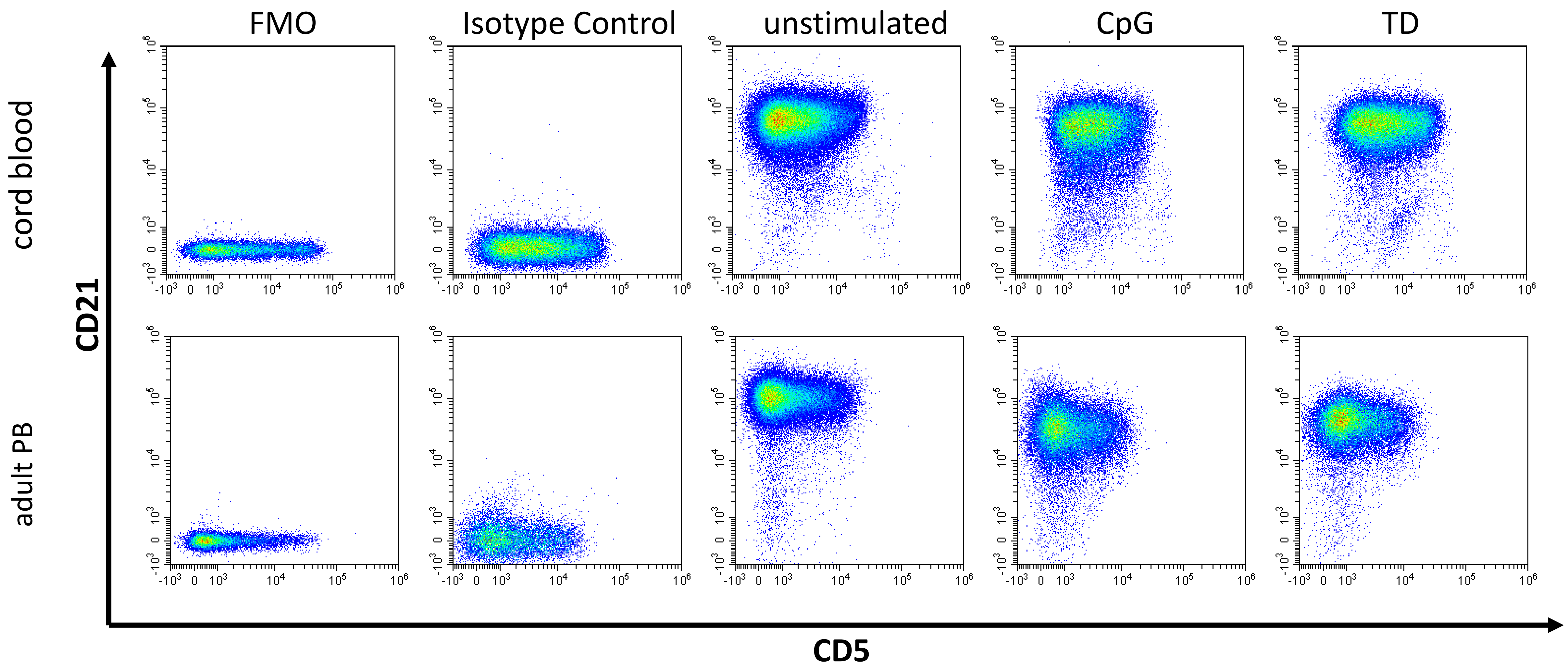

FMO
Isotype Control
unstimulated
CpG
TD
cord blood
adult PB
CD21
CD5

H

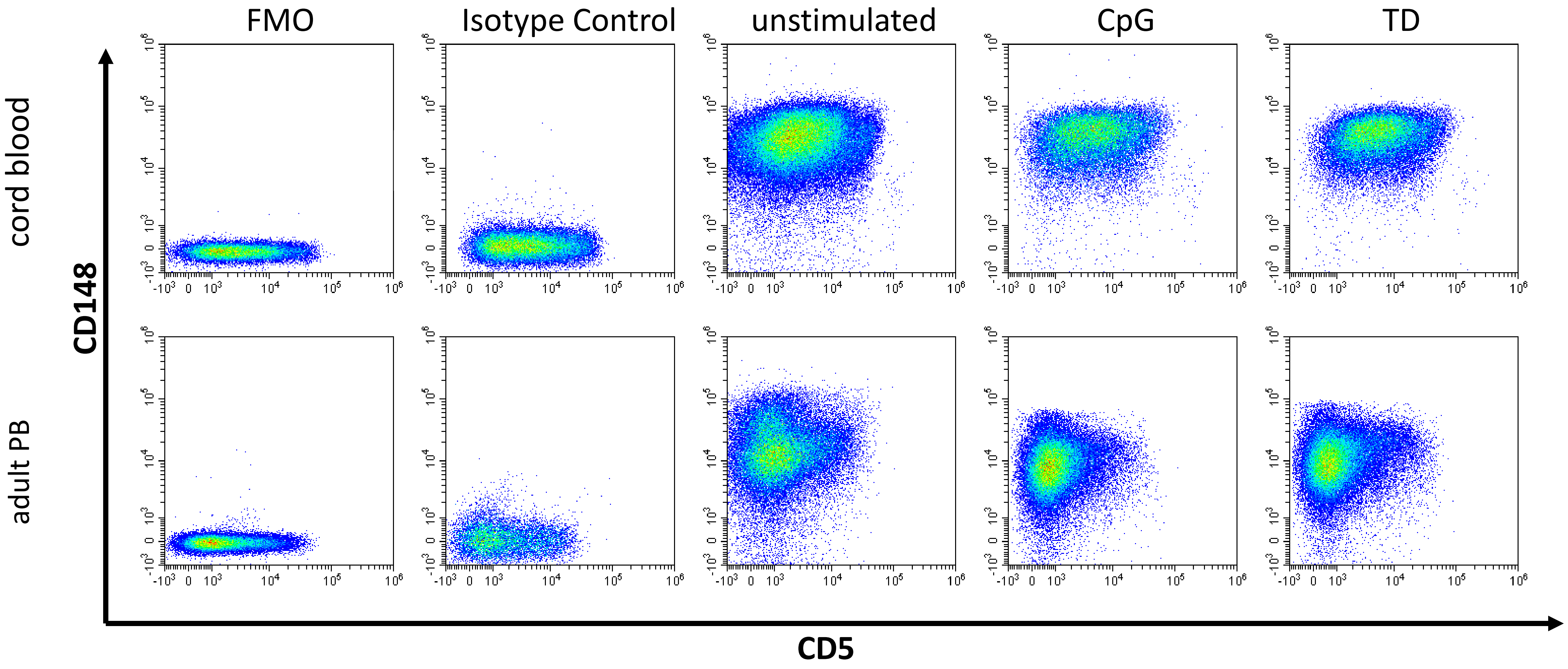

FMO
Isotype Control
unstimulated
CpG
TD
cord blood
adult PB
CD148
CD5

I

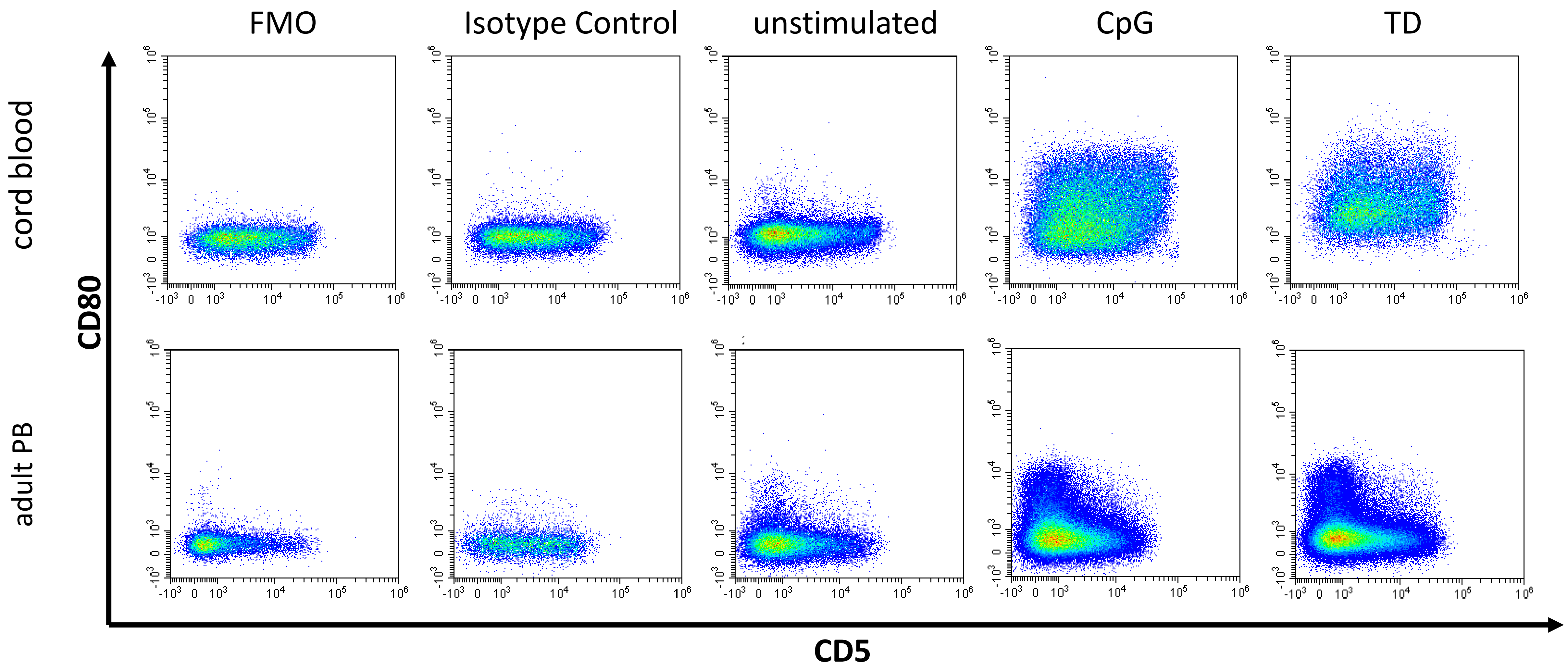
FMO
Isotype Control
unstimulated
CpG
TD
cord blood
adult PB
CD80
CD5

J

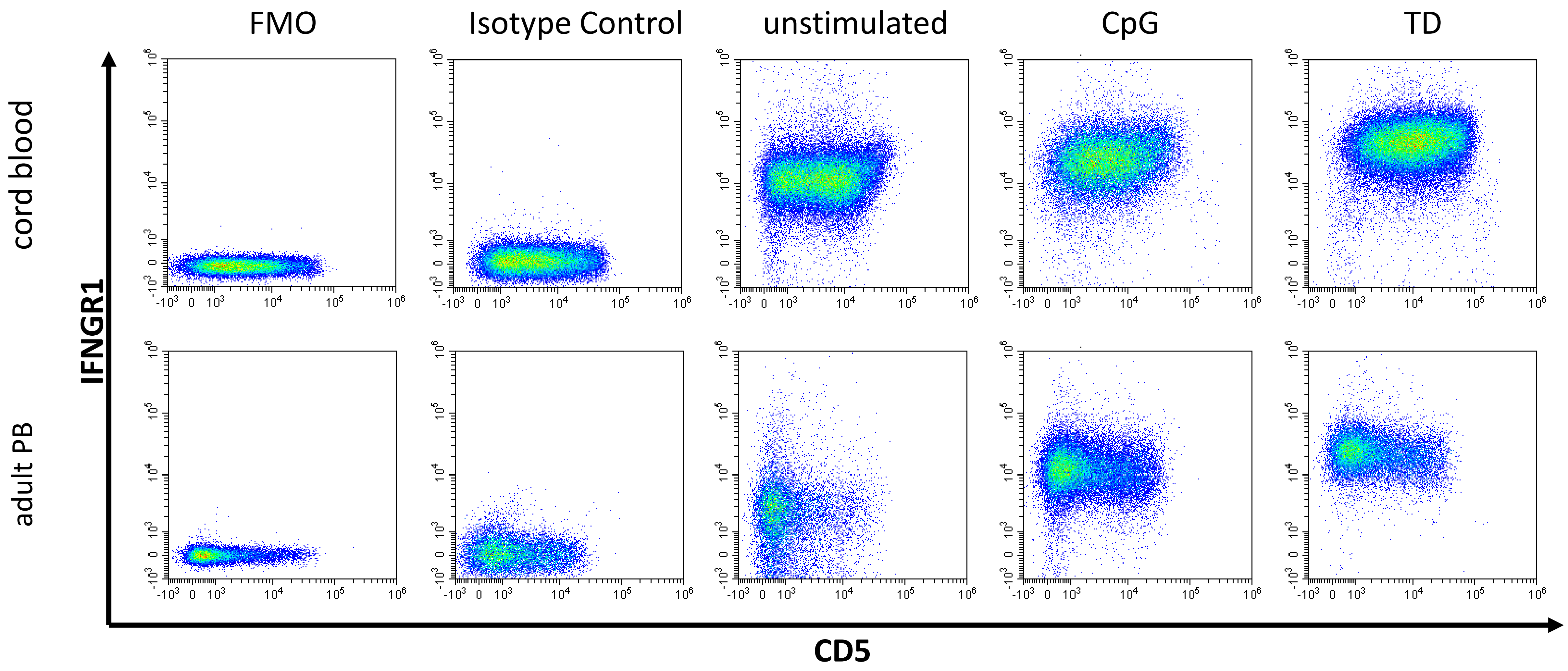

FMO
Isotype Control
unstimulated
CpG
TD
cord blood
adult PB
IFNGR1
CD5

K

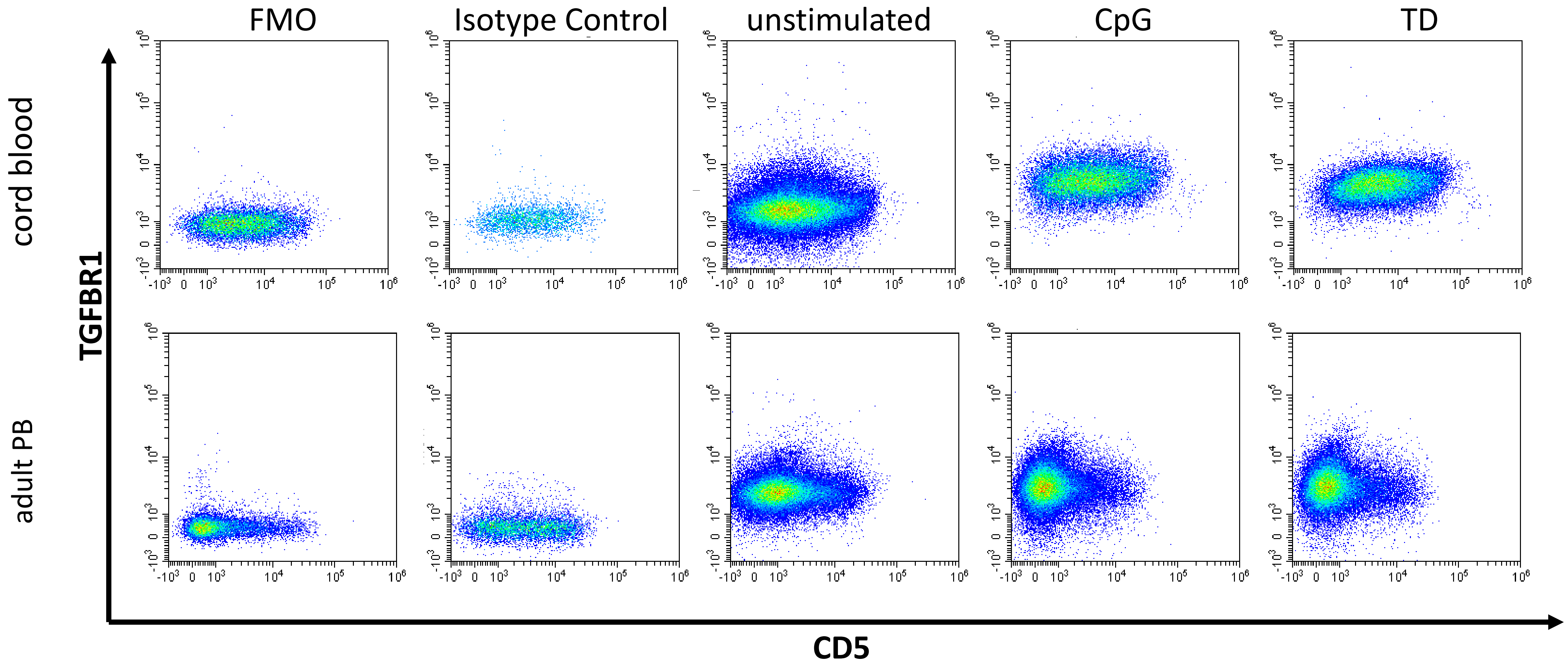
FMO
Isotype Control
unstimulated
CpG
TD
cord blood
adult PB
TGFBR1
CD5

**L**

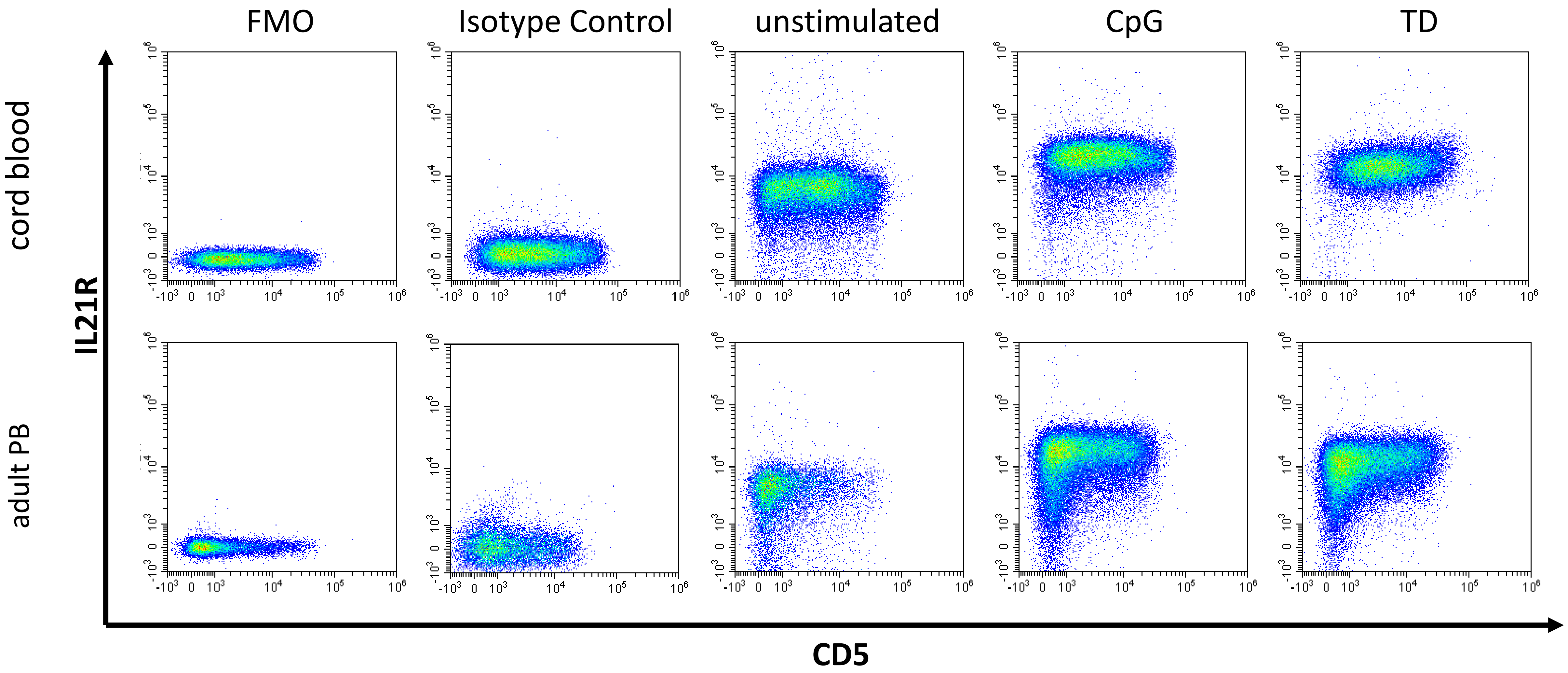
FMO
Isotype Control
unstimulated
CpG
TD
cord blood
adult PB
IL21R
CD5

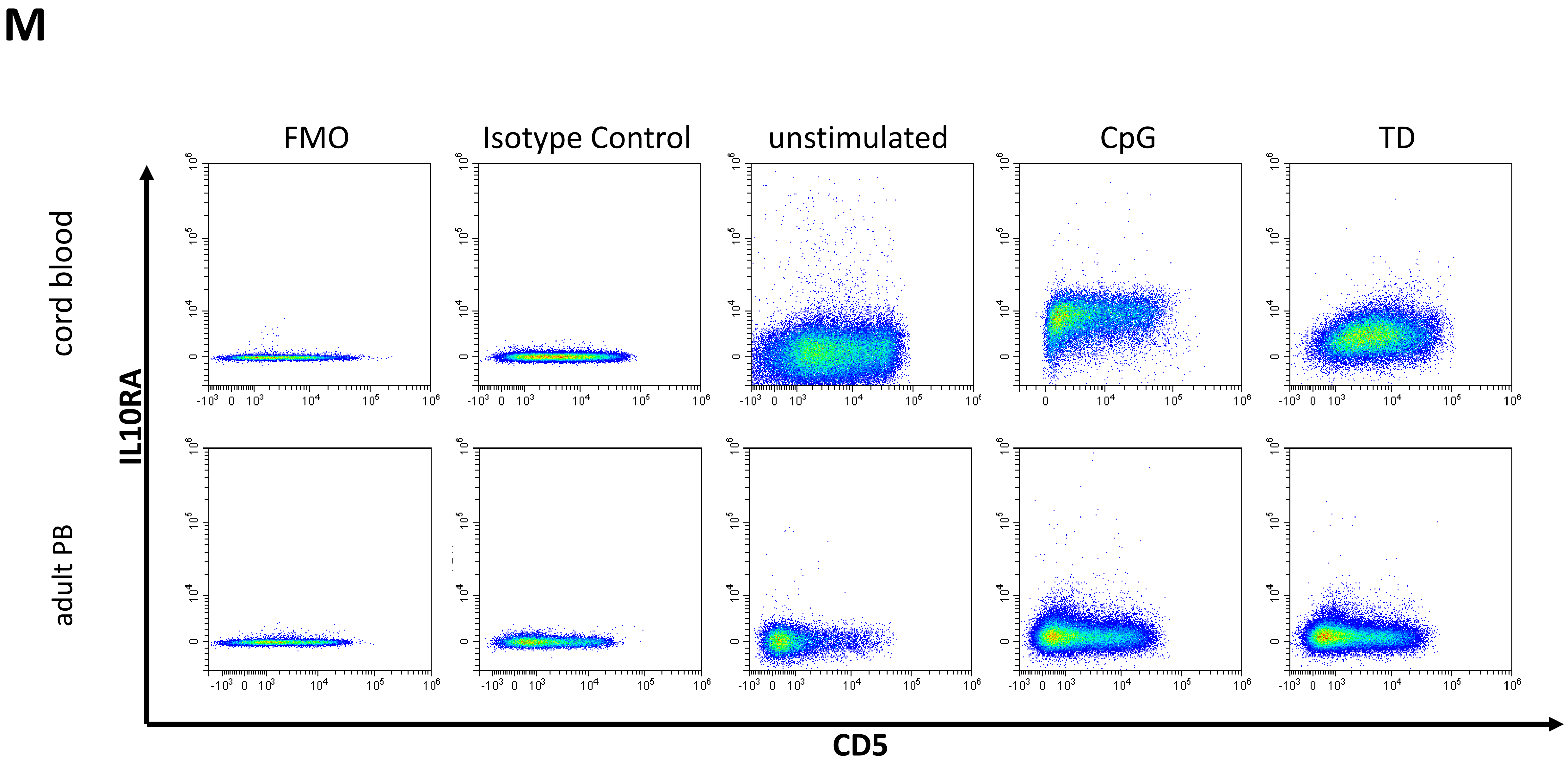
M
FMO
Isotype Control
unstimulated
CpG
TD
cord blood
adult PB
IL10RA
CD5

# N

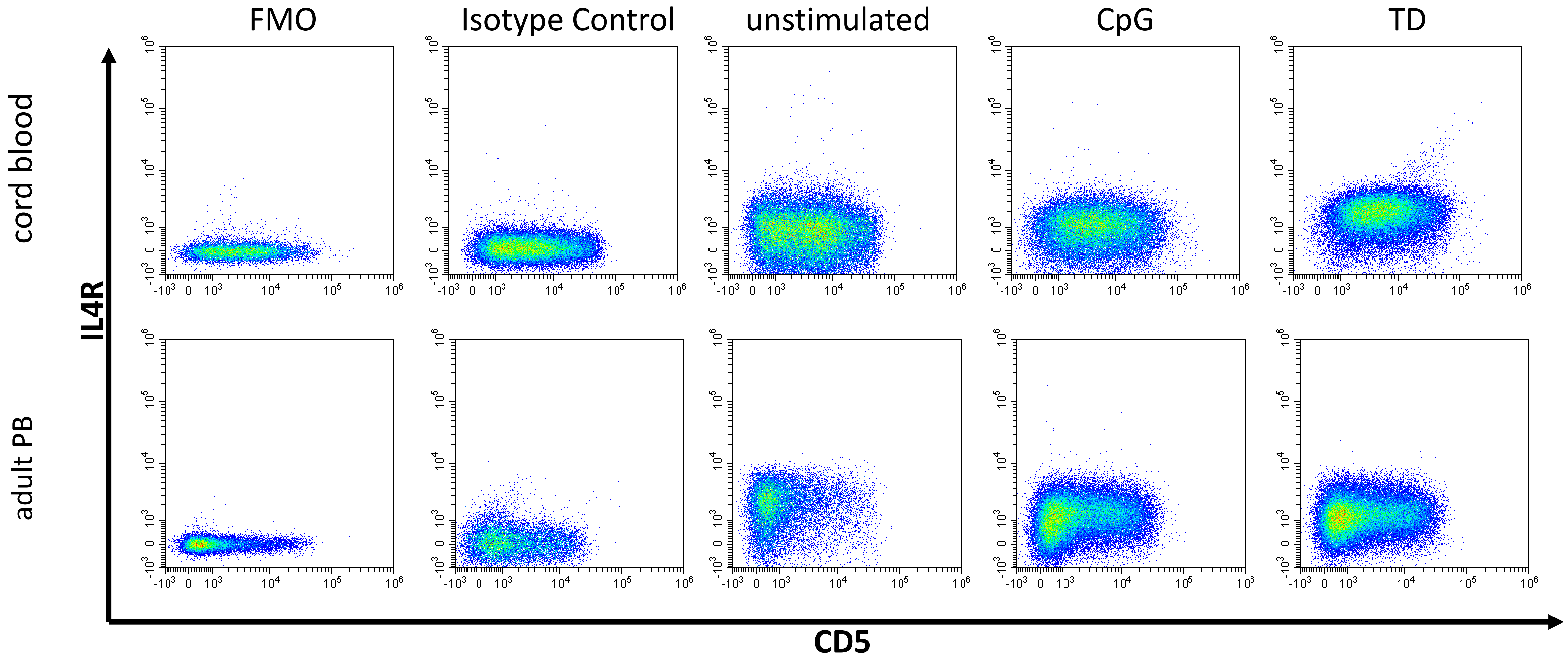
FMO
Isotype Control
unstimulated
CpG
TD
cord blood
adult PB
IL4R
CD5
