## Supplemental Figure 4 for "Human neonatal B cell immunity differs from the adult version by conserved Ig repertoires and rapid, but transient response dynamics"

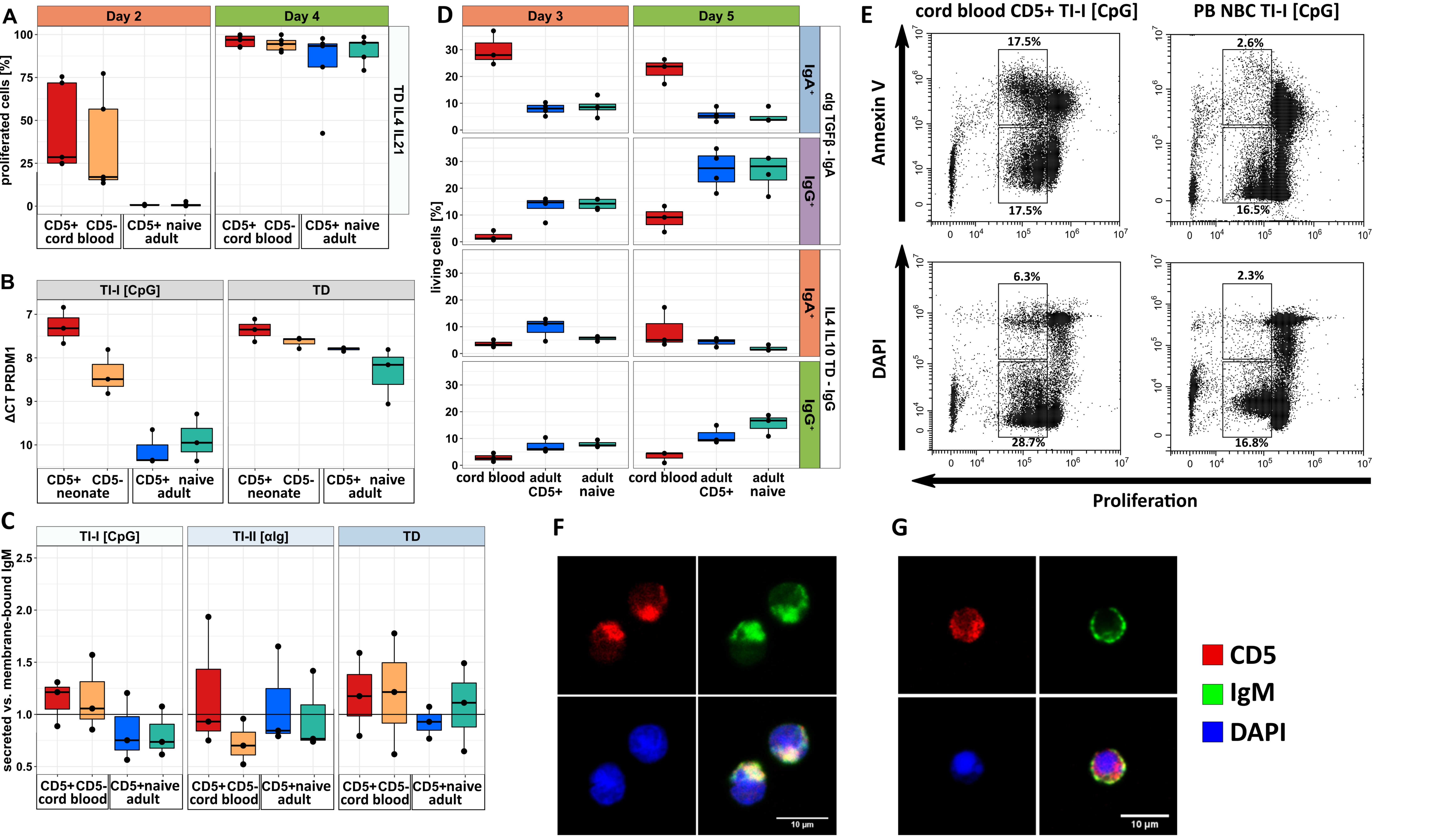


**Supplemental Figure 4, related to Figure 4. Functional Analysis of Cord Blood and Adult Mature B Cells.**

(A) Summary of five samples each of cord blood and adult mature CD5+ and CD5- B cells, proliferating upon TD plus IL-4 and IL-21 stimulation.

(B) Delta-Ct values of PRDM1 transcription upon stimulation on d3, normalized to GAPDH. PRDM1 transcription was not detectable under TI-II stimulation.

(C) Ratio of membrane-IgM and secreted-IgM transcript levels upon stimulation on d3, as determined by semi-quantitative PCR.

(D) Summary of three to five independent class switching experiments of human cord blood (CD5+ and CD5- B cells were not separated for the sake of sufficient cell numbers) and adult CD5+ and CD5- B cells, stimulated with TGFb and anti-Ig (IgA switching) or stimulated with IL-4, IL-10 and TD conditions.

(E) Apoptosis (top) and cell death (bottom) of cord blood (left) and adult (right) B cells under TI-I stimulation on d4.

(F and G) Cell polarization (IgM capping) of cord blood (F) or adult (G) mature (naïve) B cells after 30 minutes of stimulation with CpG. DAPI (blue), CD5 (red), IgM (green).
