## Supplemental Figure 5 for "Human neonatal B cell immunity differs from the adult version by conserved Ig repertoires and rapid, but transient response dynamics"

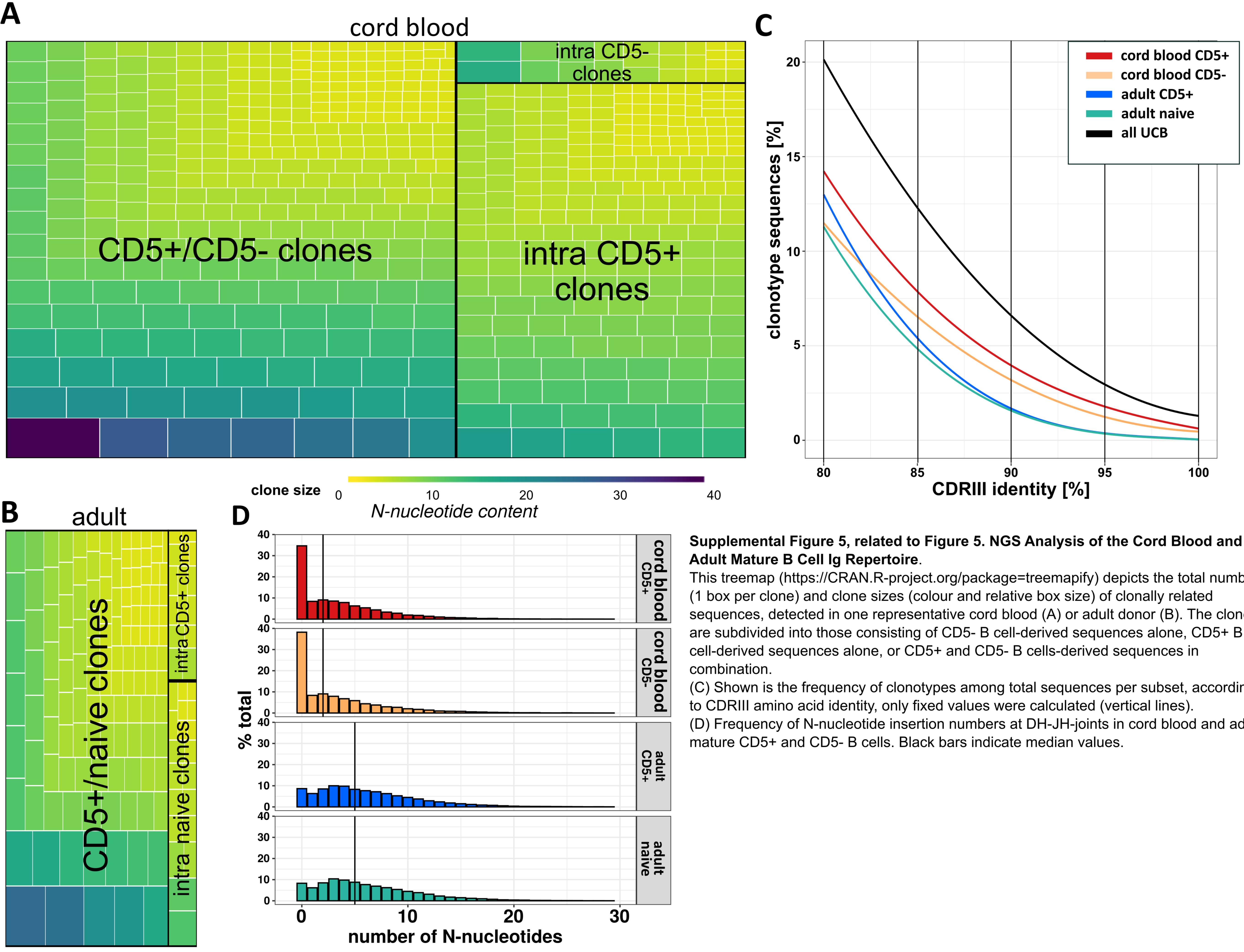


**Supplemental Figure 5, related to Figure 5. NGS Analysis of the Cord Blood and Adult Mature B Cell Ig Repertoire**.
This treemap (https://CRAN.R-project.org/package=treemapify) depicts the total number (1 box per clone) and clone sizes (colour and relative box size) of clonally related sequences, detected in one representative cord blood (A) or adult donor (B). The clones are subdivided into those consisting of CD5- B cell-derived sequences alone, CD5+ B cell-derived sequences alone, or CD5+ and CD5- B cells-derived sequences in combination.
(C) Shown is the frequency of clonotypes among total sequences per subset, according to CDRIII amino acid identity, only fixed values were calculated (vertical lines).
(D) Frequency of N-nucleotide insertion numbers at DH-JH-joints in cord blood and adult mature CD5+ and CD5- B cells. Black bars indicate median values.
