## Supplemental Table 1 for "Human neonatal B cell immunity differs from the adult version by conserved Ig repertoires and rapid, but transient response dynamics"

| Pathway | pval | padj | NES | |
|---|---|---|---|---|
| HALLMARK_TNFA_SIGNALING_VIA_NFKB | 0.00022 | 0.00222 | 2.63139 | Hallmark |
| HALLMARK_TGF_BETA_SIGNALING | 0.00043 | 0.00318 | 1.89645 | Hallmark |
| HALLMARK_UV_RESPONSE_UP | 0.00022 | 0.00222 | 1.81247 | Hallmark |
| HALLMARK_HYPOXIA | 0.00022 | 0.00222 | 1.79452 | Hallmark |
| HALLMARK_APOPTOSIS | 0.00022 | 0.00222 | 1.75860 | Hallmark |
| HALLMARK_INFLAMMATORY_RESPONSE | 0.00044 | 0.00318 | 1.75744 | Hallmark |
| HALLMARK_MYOGENESIS | 0.00022 | 0.00222 | 1.74590 | Hallmark |
| HALLMARK_IL2_STAT5_SIGNALING | 0.00088 | 0.00465 | 1.60851 | Hallmark |
| HALLMARK_G2M_CHECKPOINT | 0.00112 | 0.00465 | 1.57672 | Hallmark |
| HALLMARK_EPITHELIAL_MESENCHYMAL_TRANSITION | 0.00110 | 0.00465 | 1.56308 | Hallmark |
| HALLMARK_MITOTIC_SPINDLE | 0.00111 | 0.00465 | 1.55857 | Hallmark |
| HALLMARK_ESTROGEN_RESPONSE_EARLY | 0.00133 | 0.00510 | 1.53603 | Hallmark |
| HALLMARK_MTORC1_SIGNALING | 0.00111 | 0.00465 | 1.51423 | Hallmark |
| HALLMARK_UNFOLDED_PROTEIN_RESPONSE | 0.00414 | 0.01149 | 1.51383 | Hallmark |
| HALLMARK_P53_PATHWAY | 0.00266 | 0.00831 | 1.47797 | Hallmark |
| HALLMARK_COMPLEMENT | 0.00288 | 0.00846 | 1.46342 | Hallmark |
| HALLMARK_ANDROGEN_RESPONSE | 0.01226 | 0.03227 | 1.45590 | Hallmark |
| HALLMARK_KRAS_SIGNALING_UP | 0.02021 | 0.04393 | 1.34735 | Hallmark |
| HALLMARK_INTERFERON_ALPHA_RESPONSE | 0.00243 | 0.00809 | -1.59515 | Hallmark |
| HALLMARK_PANCREAS_BETA_CELLS | 0.01800 | 0.04393 | -1.56682 | Hallmark |
| HALLMARK_OXIDATIVE_PHOSPHORYLATION | 0.00201 | 0.00718 | -1.47561 | Hallmark |
| DIRMEIER_LMP1_RESPONSE_EARLY | 0.00021 | 0.00491 | 2.52664 | C2 |
| NAGASHIMA_NRG1_SIGNALING_UP | 0.00022 | 0.00491 | 2.51144 | C2 |
| PHONG_TNF_TARGETS_UP | 0.00021 | 0.00491 | 2.38825 | C2 |

| | | | | |
|---|---|---|---|---|
| UDAYAKUMAR_MED1_TARGETS_DN | 0.00022 | 0.00491 | 2.35638 | C2 |
| THEILGAARD_NEUTROPHIL_AT_SKIN_WOUND_UP | 0.00022 | 0.00491 | 2.35601 | C2 |
| MITSIADES_RESPONSE_TO_APLIDIN_UP | 0.00023 | 0.00491 | 2.34329 | C2 |
| RASHI_RESPONSE_TO_IONIZING_RADIATION_2 | 0.00022 | 0.00491 | 2.31142 | C2 |
| NAGASHIMA_EGF_SIGNALING_UP | 0.00021 | 0.00491 | 2.26355 | C2 |
| LINDSTEDT_DENDRITIC_CELL_MATURATION_B | 0.00021 | 0.00491 | 2.25600 | C2 |
| CHEN_HOXA5_TARGETS_9HR_UP | 0.00022 | 0.00491 | 2.22758 | C2 |
| BASSO_CD40_SIGNALING_UP | 0.00022 | 0.00491 | 2.18331 | C2 |
| ZWANG_CLASS_3_TRANSIENTLY_INDUCED_BY_EGF | 0.00022 | 0.00491 | 2.17002 | C2 |
| RHEIN_ALL_GLUCOCORTICOID_THERAPY_UP | 0.00022 | 0.00491 | 2.14579 | C2 |
| BURTON_ADIPOGENESIS_PEAK_AT_2HR | 0.00021 | 0.00491 | 2.13185 | C2 |
| ZHANG_RESPONSE_TO_IKK_INHIBITOR_AND_TNF_UP | 0.00022 | 0.00491 | 2.13014 | C2 |
| OSWALD_HEMATOPOIETIC_STEM_CELL_IN_COLLAGEN_GEL_UP | 0.00022 | 0.00491 | 2.12777 | C2 |
| UZONYI_RESPONSE_TO_LEUKOTRIENE_AND_THROMBIN | 0.00022 | 0.00491 | 2.12501 | C2 |
| PHONG_TNF_RESPONSE_NOT_VIA_P38 | 0.00023 | 0.00491 | 2.11960 | C2 |
| LEONARD_HYPOXIA | 0.00022 | 0.00491 | 2.10076 | C2 |
| DIRMEIER_LMP1_RESPONSE_LATE_UP | 0.00022 | 0.00491 | 2.09988 | C2 |
| TAKEDA_TARGETS_OF_NUP98_HOXA9_FUSION_6HR_DN | 0.00022 | 0.00491 | 2.08037 | C2 |
| AMIT_EGF_RESPONSE_120_HELA | 0.00022 | 0.00491 | 2.07730 | C2 |
| GARY_CD5_TARGETS_UP | 0.00023 | 0.00491 | 2.07571 | C2 |
| GALINDO_IMMUNE_RESPONSE_TO_ENTEROTOXIN | 0.00022 | 0.00491 | 2.07521 | C2 |
| ZHOU_INFLAMMATORY_RESPONSE_LIVE_UP | 0.00023 | 0.00491 | 2.07131 | C2 |
| VILIMAS_NOTCH1_TARGETS_UP | 0.00021 | 0.00491 | 2.06235 | C2 |
| GARGALOVIC_RESPONSE_TO_OXIDIZED_PHOSPHOLIPIDS_MAGENTA_UP | 0.00021 | 0.00491 | 2.05513 | C2 |
| TIAN_TNF_SIGNALING_VIA_NFKB | 0.00022 | 0.00491 | 2.04031 | C2 |
| KIM_WT1_TARGETS_UP | 0.00022 | 0.00491 | 2.02787 | C2 |

| | | | | |
|---|---|---|---|---|
| TIEN_INTESTINE_PROBIOTICS_24HR_DN | 0.00022 | 0.00491 | 2.02610 | C2 |
| PRAMOONJAGO_SOX4_TARGETS_UP | 0.00021 | 0.00491 | 2.01412 | C2 |
| SMITH_TERT_TARGETS_UP | 0.00022 | 0.00491 | 2.00012 | C2 |
| GARGALOVIC_RESPONSE_TO_OXIDIZED_PHOSPHOLIPIDS_BLUE_UP | 0.00022 | 0.00491 | 1.99273 | C2 |
| PID_LYSOPHOSPHOLIPID_PATHWAY | 0.00022 | 0.00491 | 1.98180 | C2 |
| RASHI_NFKB1_TARGETS | 0.00022 | 0.00491 | 1.97851 | C2 |
| AMIT_SERUM_RESPONSE_40_MCF10A | 0.00021 | 0.00491 | 1.97678 | C2 |
| CHO_NR4A1_TARGETS | 0.00043 | 0.00771 | 1.96388 | C2 |
| BENPORATH_ES_CORE_NINE_CORRELATED | 0.00022 | 0.00491 | 1.96290 | C2 |
| SEKI_INFLAMMATORY_RESPONSE_LPS_UP | 0.00022 | 0.00491 | 1.95020 | C2 |
| SMIRNOV_RESPONSE_TO_IR_2HR_UP | 0.00021 | 0.00491 | 1.94699 | C2 |
| FERRARI_RESPONSE_TO_FENRETINIDE_UP | 0.00022 | 0.00491 | 1.94044 | C2 |
| BROCKE_APOPTOSIS_REVERSED_BY_IL6 | 0.00022 | 0.00491 | 1.93382 | C2 |
| GARGALOVIC_RESPONSE_TO_OXIDIZED_PHOSPHOLIPIDS_TURQUOISE_UP | 0.00022 | 0.00491 | 1.92780 | C2 |
| ONO_FOXP3_TARGETS_UP | 0.00022 | 0.00491 | 1.92585 | C2 |
| SHAFFER_IRF4_TARGETS_IN_ACTIVATED_B_LYMPHOCYTE | 0.00022 | 0.00491 | 1.91929 | C2 |
| SCIAN_INVERSED_TARGETS_OF_TP53_AND_TP73_DN | 0.00021 | 0.00491 | 1.91910 | C2 |
| DAUER_STAT3_TARGETS_UP | 0.00022 | 0.00491 | 1.91733 | C2 |
| BIOCARTA_TNFR2_PATHWAY | 0.00022 | 0.00491 | 1.91509 | C2 |
| SHAFFER_IRF4_TARGETS_IN_PLASMA_CELL_VS_MATURE_B_LYMPHOCYTE | 0.00022 | 0.00491 | 1.91234 | C2 |
| HOEBEKE_LYMPHOID_STEM_CELL_UP | 0.00022 | 0.00491 | 1.89876 | C2 |
| TIAN_TNF_SIGNALING_NOT_VIA_NFKB | 0.00022 | 0.00491 | 1.89097 | C2 |
| HARRIS_HYPOXIA | 0.00022 | 0.00491 | 1.88866 | C2 |
| BURTON_ADIPOGENESIS_1 | 0.00021 | 0.00491 | 1.88495 | C2 |
| PID_AP1_PATHWAY | 0.00022 | 0.00491 | 1.88200 | C2 |
| GESERICK_TERT_TARGETS_DN | 0.00022 | 0.00491 | 1.87843 | C2 |

| | | | | |
|---|---|---|---|---|
| NOJIMA_SFRP2_TARGETS_UP | 0.00021 | 0.00491 | 1.87527 | C2 |
| REACTOME_G_ALPHA_Q_SIGNALLING_EVENTS | 0.00022 | 0.00491 | 1.87290 | C2 |
| DALESSIO_TSA_RESPONSE | 0.00065 | 0.01017 | 1.87031 | C2 |
| CAFFAREL_RESPONSE_TO_THC_UP | 0.00022 | 0.00491 | 1.86965 | C2 |
| REACTOME_THROMBIN_SIGNALLING_THROUGH_PROTEINASE_ACTIVATED_RECEPTORS_PARS | 0.00043 | 0.00771 | 1.85614 | C2 |
| PID_THROMBIN_PAR1_PATHWAY | 0.00022 | 0.00491 | 1.85570 | C2 |
| MAHADEVAN_RESPONSE_TO_MP470_DN | 0.00131 | 0.01546 | 1.85545 | C2 |
| PID_CD40_PATHWAY | 0.00021 | 0.00491 | 1.85311 | C2 |
| MA_MYELOID_DIFFERENTIATION_DN | 0.00043 | 0.00771 | 1.85129 | C2 |
| SIG_CD40PATHWAYMAP | 0.00043 | 0.00771 | 1.85011 | C2 |
| PID_NFKAPPAB_CANONICAL_PATHWAY | 0.00043 | 0.00771 | 1.84922 | C2 |
| RODRIGUES_NTN1_AND_DCC_TARGETS | 0.00021 | 0.00491 | 1.84879 | C2 |
| PHONG_TNF_RESPONSE_VIA_P38_PARTIAL | 0.00022 | 0.00491 | 1.84656 | C2 |
| BIOCARTA_RAS_PATHWAY | 0.00043 | 0.00771 | 1.83823 | C2 |
| ZHOU_INFLAMMATORY_RESPONSE_FIMA_UP | 0.00023 | 0.00491 | 1.83567 | C2 |
| FOSTER_TOLERANT_MACROPHAGE_DN | 0.00023 | 0.00491 | 1.83553 | C2 |
| VARELA_ZMPSTE24_TARGETS_UP | 0.00043 | 0.00771 | 1.83100 | C2 |
| PURBEY_TARGETS_OF_CTBP1_AND_SATB1_UP | 0.00022 | 0.00491 | 1.81808 | C2 |
| PID_IL8_CXCR1_PATHWAY | 0.00086 | 0.01201 | 1.81491 | C2 |
| EPPERT_HSC_R | 0.00022 | 0.00491 | 1.81064 | C2 |
| NGUYEN_NOTCH1_TARGETS_DN | 0.00022 | 0.00491 | 1.80304 | C2 |
| REACTOME_CLASS_B_2_SECRETIN_FAMILY_RECEPTORS | 0.00065 | 0.01017 | 1.80049 | C2 |
| NAKAYAMA_FRA2_TARGETS | 0.00130 | 0.01546 | 1.79783 | C2 |
| DACOSTA_UV_RESPONSE_VIA_ERCC3_COMMON_DN | 0.00023 | 0.00491 | 1.79752 | C2 |
| GRAHAM_CML_QUIESCENT_VS_NORMAL_DIVIDING_UP | 0.00129 | 0.01546 | 1.79428 | C2 |
| BILANGES_SERUM_RESPONSE_TRANSLATION | 0.00086 | 0.01201 | 1.79352 | C2 |

| | | | | |
|---|---|---|---|---|
| REACTOME_G_ALPHA_Z_SIGNALLING_EVENTS | 0.00130 | 0.01546 | 1.79315 | C2 |
| ADDYA_ERYTHROID_DIFFERENTIATION_BY_HEMIN | 0.00044 | 0.00771 | 1.79270 | C2 |
| MARTENS_BOUND_BY_PML_RARA_FUSION | 0.00023 | 0.00491 | 1.79058 | C2 |
| BIOCARTA_CXCR4_PATHWAY | 0.00130 | 0.01546 | 1.78472 | C2 |
| PID_TXA2PATHWAY | 0.00064 | 0.01017 | 1.78349 | C2 |
| JIANG_HYPOXIA_NORMAL | 0.00023 | 0.00491 | 1.78317 | C2 |
| PID_FRA_PATHWAY | 0.00107 | 0.01415 | 1.77946 | C2 |
| PID_HDAC_CLASSI_PATHWAY | 0.00044 | 0.00771 | 1.77441 | C2 |
| ST_TUMOR_NECROSIS_FACTOR_PATHWAY | 0.00107 | 0.01415 | 1.76995 | C2 |
| BIOCARTA_AKT_PATHWAY | 0.00195 | 0.01971 | 1.76493 | C2 |
| REACTOME_G_BETA_GAMMA_SIGNALLING_THROUGH_PI3KGAMMA | 0.00174 | 0.01866 | 1.76477 | C2 |
| ST_FAS_SIGNALING_PATHWAY | 0.00086 | 0.01201 | 1.76235 | C2 |
| KRIEG_HYPOXIA_VIA_KDM3A | 0.00128 | 0.01546 | 1.75973 | C2 |
| REACTOME_G_ALPHA_S_SIGNALLING_EVENTS | 0.00022 | 0.00491 | 1.75962 | C2 |
| REACTOME_PROCESSING_OF_CAPPED_INTRON_CONTAINING_PRE_MRNA | 0.00022 | 0.00491 | 1.75789 | C2 |
| GAURNIER_PSMD4_TARGETS | 0.00086 | 0.01201 | 1.75636 | C2 |
| DORSEY_GAB2_TARGETS | 0.00151 | 0.01708 | 1.75470 | C2 |
| SAFFORD_T_LYMPHOCYTE_ANERGY | 0.00022 | 0.00491 | 1.75468 | C2 |
| ST_ERK1_ERK2_MAPK_PATHWAY | 0.00129 | 0.01546 | 1.74835 | C2 |
| REACTOME_G_PROTEIN_BETA_GAMMA_SIGNALLING | 0.00151 | 0.01708 | 1.74807 | C2 |
| SUH_COEXPRESSED_WITH_ID1_AND_ID2_UP | 0.00327 | 0.02731 | 1.74648 | C2 |
| BIOCARTA_CDMAC_PATHWAY | 0.00372 | 0.02992 | 1.74512 | C2 |
| KIM_GERMINAL_CENTER_T_HELPER_DN | 0.00239 | 0.02243 | 1.74387 | C2 |
| PENG_LEUCINE_DEPRIVATION_DN | 0.00022 | 0.00491 | 1.74176 | C2 |
| SENESE_HDAC3_TARGETS_UP | 0.00023 | 0.00491 | 1.74086 | C2 |
| REACTOME_GASTRIN_CREB_SIGNALLING_PATHWAY_VIA_PKC_AND_MAPK | 0.00022 | 0.00491 | 1.74054 | C2 |

| | | | | |
|---|---|---|---|---|
| MOREAUX_B_LYMPHOCYTE_MATURATION_BY_TACI_UP | 0.00088 | 0.01201 | 1.73914 | C2 |
| REACTOME_PLATELET_ACTIVATION_SIGNALING_AND_AGGREGATION | 0.00022 | 0.00491 | 1.73776 | C2 |
| KEGG_MAPK_SIGNALING_PATHWAY | 0.00022 | 0.00491 | 1.73740 | C2 |
| MARCHINI_TRABECTEDIN_RESISTANCE_DN | 0.00151 | 0.01708 | 1.73651 | C2 |
| ZHOU_TNF_SIGNALING_30MIN | 0.00171 | 0.01865 | 1.73328 | C2 |
| KRIGE_AMINO_ACID_DEPRIVATION | 0.00215 | 0.02083 | 1.73179 | C2 |
| ZHOU_INFLAMMATORY_RESPONSE_LPS_UP | 0.00023 | 0.00491 | 1.72892 | C2 |
| ZWANG_CLASS_1_TRANSIENTLY_INDUCED_BY_EGF | 0.00023 | 0.00491 | 1.72581 | C2 |
| XU_HGF_SIGNALING_NOT_VIA_AKT1_6HR | 0.00237 | 0.02236 | 1.72520 | C2 |
| REACTOME_STRIATED_MUSCLE_CONTRACTION | 0.00522 | 0.03783 | 1.72489 | C2 |
| HINATA_NFKB_TARGETS_KERATINOCYTE_UP | 0.00088 | 0.01201 | 1.72441 | C2 |
| BAKER_HEMATOPOIESIS_STAT3_TARGETS | 0.00436 | 0.03363 | 1.72386 | C2 |
| YIH_RESPONSE_TO_ARSENITE_C1 | 0.00326 | 0.02731 | 1.72333 | C2 |
| REACTOME_THROMBOXANE_SIGNALLING_THROUGH_TP_RECEPTOR | 0.00369 | 0.02981 | 1.72130 | C2 |
| KEGG_PATHWAYS_IN_CANCER | 0.00023 | 0.00491 | 1.71910 | C2 |
| LAU_APOPTOSIS_CDKN2A_UP | 0.00172 | 0.01866 | 1.71613 | C2 |
| BOQUEST_STEM_CELL_CULTURED_VS_FRESH_UP | 0.00023 | 0.00491 | 1.71214 | C2 |
| NATSUME_RESPONSE_TO_INTERFERON_BETA_UP | 0.00108 | 0.01415 | 1.71131 | C2 |
| PID_NFKAPPAB_ATYPICAL_PATHWAY | 0.00588 | 0.04109 | 1.71075 | C2 |
| ZHONG_RESPONSE_TO_AZACITIDINE_AND_TSA_UP | 0.00022 | 0.00491 | 1.70651 | C2 |
| KEGG_NEUROTROPHIN_SIGNALING_PATHWAY | 0.00088 | 0.01201 | 1.70456 | C2 |
| ST_B_CELL_ANTIGEN_RECEPTOR | 0.00284 | 0.02511 | 1.70412 | C2 |
| AMUNDSON_GENOTOXIC_SIGNATURE | 0.00044 | 0.00771 | 1.69940 | C2 |
| PARK_HSC_MARKERS | 0.00342 | 0.02803 | 1.69842 | C2 |
| KEGG_ERBB_SIGNALING_PATHWAY | 0.00044 | 0.00771 | 1.69814 | C2 |
| ZHANG_ANTIVIRAL_RESPONSE_TO_RIBAVIRIN_DN | 0.00324 | 0.02731 | 1.69798 | C2 |

| | | | | |
|---|---|---|---|---|
| REACTOME_SEMAPHORIN_INTERACTIONS | 0.00129 | 0.01546 | 1.69731 | C2 |
| NEMETH_INFLAMMATORY_RESPONSE_LPS_UP | 0.00044 | 0.00771 | 1.69554 | C2 |
| YAMAZAKI_TCEB3_TARGETS_UP | 0.00088 | 0.01201 | 1.69212 | C2 |
| WELCSH_BRCA1_TARGETS_DN | 0.00088 | 0.01201 | 1.68855 | C2 |
| REACTOME_TRANSPORT_OF_MATURE_TRANSCRIPT_TO_CYTOPLASM | 0.00257 | 0.02367 | 1.68855 | C2 |
| TENEDINI_MEGAKARYOCYTE_MARKERS | 0.00129 | 0.01546 | 1.68836 | C2 |
| HSIAO_HOUSEKEEPING_GENES | 0.00023 | 0.00491 | 1.68781 | C2 |
| WINTER_HYPOXIA_METAGENE | 0.00022 | 0.00491 | 1.68336 | C2 |
| SASSON_RESPONSE_TO_FORSKOLIN_UP | 0.00066 | 0.01017 | 1.68280 | C2 |
| LEI_MYB_TARGETS | 0.00023 | 0.00491 | 1.68186 | C2 |
| HELLER_HDAC_TARGETS_SILENCED_BY_METHYLATION_DN | 0.00022 | 0.00491 | 1.68135 | C2 |
| KIM_WT1_TARGETS_8HR_UP | 0.00088 | 0.01201 | 1.68010 | C2 |
| PHONG_TNF_RESPONSE_VIA_P38_COMPLETE | 0.00022 | 0.00491 | 1.67944 | C2 |
| LOPEZ_TRANSLATION_VIA_FN1_SIGNALING | 0.00516 | 0.03767 | 1.67905 | C2 |
| JACKSON_DNMT1_TARGETS_UP | 0.00173 | 0.01866 | 1.67730 | C2 |
| ODONNELL_TFRC_TARGETS_UP | 0.00023 | 0.00491 | 1.67178 | C2 |
| CROONQUIST_STROMAL_STIMULATION_UP | 0.00279 | 0.02507 | 1.67088 | C2 |
| PID_ERBB1_INTERNALIZATION_PATHWAY | 0.00476 | 0.03563 | 1.66975 | C2 |
| PENG_GLUTAMINE_DEPRIVATION_DN | 0.00023 | 0.00491 | 1.66909 | C2 |
| GROSS_HYPOXIA_VIA_ELK3_DN | 0.00088 | 0.01201 | 1.66625 | C2 |
| PID_MET_PATHWAY | 0.00066 | 0.01017 | 1.66388 | C2 |
| PEREZ_TP63_TARGETS | 0.00022 | 0.00491 | 1.66326 | C2 |
| WATTEL_AUTONOMOUS_THYROID_ADENOMA_DN | 0.00409 | 0.03208 | 1.66321 | C2 |
| REACTOME_GPVI_MEDIATED_ACTIVATION_CASCADE | 0.00559 | 0.03969 | 1.66304 | C2 |
| BIOCARTA_RACCYCD_PATHWAY | 0.00662 | 0.04398 | 1.66089 | C2 |
| SARRIO_EPITHELIAL_MESENCHYMAL_TRANSITION_DN | 0.00088 | 0.01201 | 1.66015 | C2 |

| | | | | |
|---|---|---|---|---|
| BIOCARTA_PAR1_PATHWAY | 0.00502 | 0.03708 | 1.65886 | C2 |
| PID_IGF1_PATHWAY | 0.00598 | 0.04145 | 1.65828 | C2 |
| MILI_PSEUDOPODIA_CHEMOTAXIS_DN | 0.00023 | 0.00491 | 1.65404 | C2 |
| GRAHAM_NORMAL_QUIESCENT_VS_NORMAL_DIVIDING_UP | 0.00216 | 0.02083 | 1.65358 | C2 |
| REACTOME_MRNA_SPLICING | 0.00088 | 0.01201 | 1.65200 | C2 |
| MORI_IMMATURE_B_LYMPHOCYTE_UP | 0.00386 | 0.03064 | 1.65012 | C2 |
| UEDA_CENTRAL_CLOCK | 0.00066 | 0.01017 | 1.64959 | C2 |
| REACTOME_GLUCAGON_TYPE_LIGAND_RECEPTORS | 0.00734 | 0.04637 | 1.64891 | C2 |
| ST_JNK_MAPK_PATHWAY | 0.00541 | 0.03887 | 1.64843 | C2 |
| REACTOME_SIGNAL_AMPLIFICATION | 0.00809 | 0.04973 | 1.64670 | C2 |
| PID_IL8_CXCR2_PATHWAY | 0.00517 | 0.03769 | 1.64554 | C2 |
| SHEPARD_CRUSH_AND_BURN_MUTANT_DN | 0.00110 | 0.01415 | 1.64515 | C2 |
| ST_DIFFERENTIATION_PATHWAY_IN_PC12_CELLS | 0.00541 | 0.03887 | 1.64510 | C2 |
| MENSE_HYPOXIA_UP | 0.00066 | 0.01017 | 1.64145 | C2 |
| ELVIDGE_HYPOXIA_UP | 0.00088 | 0.01201 | 1.64135 | C2 |
| KANNAN_TP53_TARGETS_UP | 0.00410 | 0.03208 | 1.64118 | C2 |
| PID_AMB2_NEUTROPHILS_PATHWAY | 0.00611 | 0.04198 | 1.63937 | C2 |
| PID_HIF1_TFPATHWAY | 0.00260 | 0.02380 | 1.63794 | C2 |
| PENG_RAPAMYCIN_RESPONSE_DN | 0.00022 | 0.00491 | 1.63714 | C2 |
| BIOCARTA_MAPK_PATHWAY | 0.00089 | 0.01201 | 1.63668 | C2 |
| PID_INSULIN_PATHWAY | 0.00470 | 0.03557 | 1.63402 | C2 |
| LU_AGING_BRAIN_DN | 0.00132 | 0.01546 | 1.63267 | C2 |
| SIG_PIP3_SIGNALING_IN_CARDIAC_MYOCTES | 0.00260 | 0.02380 | 1.62802 | C2 |
| DACOSTA_UV_RESPONSE_VIA_ERCC3_COMMON_UP | 0.00218 | 0.02101 | 1.62704 | C2 |
| PLASARI_TGFB1_TARGETS_10HR_UP | 0.00088 | 0.01201 | 1.62607 | C2 |
| HINATA_NFKB_TARGETS_FIBROBLAST_UP | 0.00240 | 0.02244 | 1.62522 | C2 |

| | | | | |
|---|---|---|---|---|
| ZHANG_TARGETS_OF_EWSR1_FLI1_FUSION | 0.00176 | 0.01866 | 1.62179 | C2 |
| PID_ILK_PATHWAY | 0.00513 | 0.03764 | 1.62161 | C2 |
| RUTELLA_RESPONSE_TO_HGF_VS_CSF2RB_AND_IL4_DN | 0.00022 | 0.00491 | 1.62122 | C2 |
| DELPUECH_FOXO3_TARGETS_UP | 0.00345 | 0.02820 | 1.61924 | C2 |
| SCHEIDEREIT_IKK_INTERACTING_PROTEINS | 0.00302 | 0.02607 | 1.61657 | C2 |
| ZHANG_TLX_TARGETS_UP | 0.00132 | 0.01546 | 1.61133 | C2 |
| YAO_TEMPORAL_RESPONSE_TO_PROGESTERONE_CLUSTER_7 | 0.00306 | 0.02627 | 1.60436 | C2 |
| REACTOME_DOWNSTREAM_SIGNALING_EVENTS_OF_B_CELL_RECEPTOR_BCR | 0.00132 | 0.01546 | 1.60311 | C2 |
| QI_HYPOXIA | 0.00132 | 0.01546 | 1.60276 | C2 |
| WIERENGA_STAT5A_TARGETS_GROUP2 | 0.00600 | 0.04152 | 1.59983 | C2 |
| REACTOME_3_UTR_MEDIATED_TRANSLATIONAL_REGULATION | 0.00132 | 0.01546 | 1.59897 | C2 |
| MARTORIATI_MDM4_TARGETS_FETAL_LIVER_DN | 0.00023 | 0.00491 | 1.59869 | C2 |
| PENG_LEUCINE_DEPRIVATION_UP | 0.00132 | 0.01546 | 1.59649 | C2 |
| MANALO_HYPOXIA_UP | 0.00044 | 0.00771 | 1.59582 | C2 |
| CROONQUIST_NRAS_VS_STROMAL_STIMULATION_DN | 0.00110 | 0.01415 | 1.59420 | C2 |
| MEISSNER_BRAIN_HCP_WITH_H3K27ME3 | 0.00154 | 0.01727 | 1.59398 | C2 |
| REACTOME_ACTIVATION_OF_THE_MRNA_UPON_BINDING_OF_THE_CAP_BINDING_COMPLEX_AND_EIFS_AND_SUBSEQUENT_BINDING_TO_43S | 0.00412 | 0.03220 | 1.59233 | C2 |
| IVANOVA_HEMATOPOIESIS_STEM_CELL | 0.00022 | 0.00491 | 1.59078 | C2 |
| GROSS_HYPOXIA_VIA_ELK3_AND_HIF1A_UP | 0.00132 | 0.01546 | 1.59076 | C2 |
| RUTELLA_RESPONSE_TO_HGF_DN | 0.00022 | 0.00491 | 1.59021 | C2 |
| PID_TGFBR_PATHWAY | 0.00473 | 0.03563 | 1.59004 | C2 |
| KIM_WT1_TARGETS_12HR_UP | 0.00153 | 0.01717 | 1.58825 | C2 |
| EBAUER_TARGETS_OF_PAX3_FOXO1_FUSION_UP | 0.00066 | 0.01017 | 1.58153 | C2 |
| WIERENGA_STAT5A_TARGETS_UP | 0.00022 | 0.00491 | 1.58075 | C2 |
| REACTOME_MRNA_PROCESSING | 0.00132 | 0.01546 | 1.57874 | C2 |

| | | | | |
|---|---|---|---|---|
| MIKKELSEN_NPC_HCP_WITH_H3K4ME3_AND_H3K27ME3 | 0.00153 | 0.01717 | 1.57794 | C2 |
| REACTOME_SIGNALING_BY_TGF_BETA_RECEPTOR_COMPLEX | 0.00625 | 0.04238 | 1.57566 | C2 |
| PID_MYC_ACTIV_PATHWAY | 0.00374 | 0.02992 | 1.57335 | C2 |
| REACTOME_HEMOSTASIS | 0.00023 | 0.00491 | 1.57280 | C2 |
| BHAT_ESR1_TARGETS_VIA_AKT1_UP | 0.00022 | 0.00491 | 1.57240 | C2 |
| SASSON_RESPONSE_TO_GONADOTROPHINS_DN | 0.00330 | 0.02742 | 1.57138 | C2 |
| SABATES_COLORECTAL_ADENOMA_UP | 0.00647 | 0.04353 | 1.57041 | C2 |
| BRUECKNER_TARGETS_OF_MIRLET7A3_UP | 0.00267 | 0.02416 | 1.56860 | C2 |
| BENPORATH_MYC_TARGETS_WITH_EBOX | 0.00022 | 0.00491 | 1.56856 | C2 |
| REACTOME_SIGNALING_BY_ERBB2 | 0.00221 | 0.02112 | 1.56707 | C2 |
| REACTOME_UNFOLDED_PROTEIN_RESPONSE | 0.00374 | 0.02992 | 1.56420 | C2 |
| ZWANG_EGF_INTERVAL_DN | 0.00066 | 0.01017 | 1.56350 | C2 |
| SENGUPTA_EBNA1_ANTICORRELATED | 0.00198 | 0.01978 | 1.56300 | C2 |
| HELLER_SILENCED_BY_METHYLATION_DN | 0.00286 | 0.02511 | 1.56051 | C2 |
| RODRIGUES_NTN1_TARGETS_DN | 0.00175 | 0.01866 | 1.55992 | C2 |
| REACTOME_GPCR_DOWNSTREAM_SIGNALING | 0.00023 | 0.00491 | 1.55919 | C2 |
| BILANGES_SERUM_AND_RAPAMYCIN_SENSITIVE_GENES | 0.00715 | 0.04610 | 1.55543 | C2 |
| COULOUARN_TEMPORAL_TGFB1_SIGNATURE_UP | 0.00331 | 0.02742 | 1.54828 | C2 |
| BILANGES_SERUM_SENSITIVE_GENES | 0.00504 | 0.03709 | 1.54795 | C2 |
| LIU_CMYB_TARGETS_UP | 0.00175 | 0.01866 | 1.54729 | C2 |
| GERY_CEBP_TARGETS | 0.00286 | 0.02511 | 1.54681 | C2 |
| GUO_HEX_TARGETS_UP | 0.00526 | 0.03808 | 1.54495 | C2 |
| ELVIDGE_HYPOXIA_BY_DMOG_UP | 0.00306 | 0.02627 | 1.54494 | C2 |
| HILLION_HMGA1_TARGETS | 0.00551 | 0.03922 | 1.54305 | C2 |
| KEGG_FC_GAMMA_R_MEDIATED_PHAGOCYTOSIS | 0.00395 | 0.03126 | 1.54136 | C2 |
| KARLSSON_TGFB1_TARGETS_UP | 0.00197 | 0.01978 | 1.53933 | C2 |

| | | | | |
|---|---|---|---|---|
| KEGG_SPLICEOSOME | 0.00220 | 0.02112 | 1.53800 | C2 |
| WANG_LMO4_TARGETS_DN | 0.00046 | 0.00771 | 1.53786 | C2 |
| MEISSNER_NPC_HCP_WITH_H3K4ME3_AND_H3K27ME3 | 0.00419 | 0.03262 | 1.53600 | C2 |
| PID_P53_DOWNSTREAM_PATHWAY | 0.00263 | 0.02400 | 1.52642 | C2 |
| HADDAD_B_LYMPHOCYTE_PROGENITOR | 0.00067 | 0.01017 | 1.52196 | C2 |
| FISCHER_G2_M_CELL_CYCLE | 0.00045 | 0.00771 | 1.52032 | C2 |
| MULLIGHAN_NPM1_SIGNATURE_3_DN | 0.00264 | 0.02402 | 1.51904 | C2 |
| MISSIAGLIA_REGULATED_BY_METHYLATION_UP | 0.00329 | 0.02742 | 1.51818 | C2 |
| REACTOME_SIGNALING_BY_FGFR_IN_DISEASE | 0.00373 | 0.02992 | 1.51346 | C2 |
| WHITFIELD_CELL_CYCLE_G2_M | 0.00067 | 0.01017 | 1.51195 | C2 |
| NABA_CORE_MATRISOME | 0.00111 | 0.01415 | 1.50717 | C2 |
| PASINI_SUZ12_TARGETS_DN | 0.00068 | 0.01021 | 1.50615 | C2 |
| REACTOME_DEVELOPMENTAL_BIOLOGY | 0.00046 | 0.00771 | 1.50565 | C2 |
| KEGG_T_CELL_RECEPTOR_SIGNALING_PATHWAY | 0.00504 | 0.03709 | 1.50458 | C2 |
| DANG_REGULATED_BY_MYC_DN | 0.00067 | 0.01017 | 1.50374 | C2 |
| HELLER_HDAC_TARGETS_UP | 0.00045 | 0.00771 | 1.49684 | C2 |
| MCBRYAN_PUBERTAL_BREAST_5_6WK_DN | 0.00396 | 0.03126 | 1.49327 | C2 |
| PEREZ_TP53_AND_TP63_TARGETS | 0.00242 | 0.02253 | 1.49164 | C2 |
| MARZEC_IL2_SIGNALING_UP | 0.00441 | 0.03382 | 1.49122 | C2 |
| SENESE_HDAC1_TARGETS_UP | 0.00046 | 0.00771 | 1.48361 | C2 |
| TARTE_PLASMA_CELL_VS_PLASMABLAST_UP | 0.00046 | 0.00771 | 1.47848 | C2 |
| SHEN_SMARCA2_TARGETS_DN | 0.00157 | 0.01742 | 1.47593 | C2 |
| DELACROIX_RARG_BOUND_MEF | 0.00046 | 0.00771 | 1.47584 | C2 |
| FERNANDEZ_BOUND_BY_MYC | 0.00331 | 0.02742 | 1.47465 | C2 |
| REACTOME_SIGNALING_BY_THE_B_CELL_RECEPTOR_BCR | 0.00505 | 0.03709 | 1.47413 | C2 |
| ONDER_CDH1_TARGETS_1_UP | 0.00549 | 0.03921 | 1.47207 | C2 |

| | | | | |
|---|---|---|---|---|
| JOHNSTONE_PARVB_TARGETS_3_UP | 0.00046 | 0.00771 | 1.47038 | C2 |
| DUTERTRE_ESTRADIOL_RESPONSE_24HR_DN | 0.00023 | 0.00491 | 1.46937 | C2 |
| REACTOME_SIGNALING_BY_GPCR | 0.00023 | 0.00491 | 1.46652 | C2 |
| GAVIN_FOXP3_TARGETS_CLUSTER_P3 | 0.00374 | 0.02992 | 1.46347 | C2 |
| HELLER_SILENCED_BY_METHYLATION_UP | 0.00134 | 0.01562 | 1.45993 | C2 |
| OKUMURA_INFLAMMATORY_RESPONSE_LPS | 0.00285 | 0.02511 | 1.45724 | C2 |
| NABA_ECM_GLYCOPROTEINS | 0.00615 | 0.04213 | 1.45679 | C2 |
| KEGG_CHEMOKINE_SIGNALING_PATHWAY | 0.00483 | 0.03598 | 1.45070 | C2 |
| SENESE_HDAC1_AND_HDAC2_TARGETS_UP | 0.00289 | 0.02533 | 1.45052 | C2 |
| BASAKI_YBX1_TARGETS_DN | 0.00068 | 0.01025 | 1.44872 | C2 |
| HELLER_HDAC_TARGETS_DN | 0.00180 | 0.01880 | 1.44463 | C2 |
| ZHENG_BOUND_BY_FOXP3 | 0.00023 | 0.00491 | 1.44440 | C2 |
| REACTOME_GPCR_LIGAND_BINDING | 0.00201 | 0.01987 | 1.44421 | C2 |
| MARTORIATI_MDM4_TARGETS_FETAL_LIVER_UP | 0.00312 | 0.02667 | 1.43660 | C2 |
| MARTINEZ_RESPONSE_TO_TRABECTEDIN_DN | 0.00226 | 0.02160 | 1.43452 | C2 |
| DACOSTA_UV_RESPONSE_VIA_ERCC3_UP | 0.00181 | 0.01880 | 1.42642 | C2 |
| BRUINS_UVC_RESPONSE_VIA_TP53_GROUP_D | 0.00291 | 0.02534 | 1.41286 | C2 |
| SANSOM_APC_MYC_TARGETS | 0.00466 | 0.03537 | 1.40521 | C2 |
| HOSHIDA_LIVER_CANCER_SUBCLASS_S1 | 0.00403 | 0.03176 | 1.40265 | C2 |
| HAN_SATB1_TARGETS_DN | 0.00184 | 0.01880 | 1.39644 | C2 |
| DELYS_THYROID_CANCER_UP | 0.00137 | 0.01588 | 1.39499 | C2 |
| BHAT_ESR1_TARGETS_NOT_VIA_AKT1_UP | 0.00619 | 0.04213 | 1.39486 | C2 |
| OSWALD_HEMATOPOIETIC_STEM_CELL_IN_COLLAGEN_GEL_DN | 0.00381 | 0.03037 | 1.38405 | C2 |
| REACTOME_ADAPTIVE_IMMUNE_SYSTEM | 0.00141 | 0.01617 | 1.36275 | C2 |
| DOUGLAS_BMI1_TARGETS_UP | 0.00163 | 0.01798 | 1.35796 | C2 |
| RUTELLA_RESPONSE_TO_CSF2RB_AND_IL4_DN | 0.00341 | 0.02801 | 1.35158 | C2 |

| | | | | |
|---|---|---|---|---|
| WANG_RESPONSE_TO_GSK3_INHIBITOR_SB216763_DN | 0.00410 | 0.03208 | 1.33324 | C2 |
| MARTINEZ_TP53_TARGETS_DN | 0.00281 | 0.02511 | 1.33288 | C2 |
| MARTENS_TRETINOIN_RESPONSE_UP | 0.00326 | 0.02731 | 1.32913 | C2 |
| BRUINS_UVC_RESPONSE_VIA_TP53_GROUP_B | 0.00326 | 0.02731 | 1.32409 | C2 |
| SPIELMAN_LYMPHOBLAST_EUROPEAN_VS_ASIAN_UP | 0.00327 | 0.02731 | 1.32164 | C2 |
| GRYDER_PAX3FOXO1_TOP_ENHANCERS | 0.00438 | 0.03367 | 1.30923 | C2 |
| NABA_MATRISOME_ASSOCIATED | 0.00443 | 0.03390 | 1.30293 | C2 |
| MARTINEZ_RB1_AND_TP53_TARGETS_DN | 0.00607 | 0.04192 | 1.28696 | C2 |
| KEGG_PEROXISOME | 0.00018 | 0.00491 | -1.93772 | C2 |
| HECKER_IFNB1_TARGETS | 0.00018 | 0.00491 | -1.92205 | C2 |
| MOSERLE_IFNA_RESPONSE | 0.00037 | 0.00752 | -1.89132 | C2 |
| RHEIN_ALL_GLUCOCORTICOID_THERAPY_DN | 0.00018 | 0.00491 | -1.86254 | C2 |
| REACTOME_EXTENSION_OF_TELOMERES | 0.00056 | 0.00922 | -1.80293 | C2 |
| WONG_MITOCHONDRIA_GENE_MODULE | 0.00018 | 0.00491 | -1.79713 | C2 |
| DAUER_STAT3_TARGETS_DN | 0.00056 | 0.00922 | -1.77428 | C2 |
| REACTOME_LAGGING_STRAND_SYNTHESIS | 0.00166 | 0.01821 | -1.74648 | C2 |
| REACTOME_RNA_POL_I_TRANSCRIPTION_INITIATION | 0.00280 | 0.02507 | -1.72567 | C2 |
| REACTOME_TRANSCRIPTION_COUPLED_NER_TC_NER | 0.00150 | 0.01708 | -1.72008 | C2 |
| KEGG_DNA_REPLICATION | 0.00203 | 0.01993 | -1.71550 | C2 |
| MOOTHA_MITOCHONDRIA | 0.00018 | 0.00491 | -1.71360 | C2 |
| MOOTHA_HUMAN_MITODB_6_2002 | 0.00018 | 0.00491 | -1.71281 | C2 |
| KIM_MYCL1_AMPLIFICATION_TARGETS_DN | 0.00295 | 0.02568 | -1.69781 | C2 |
| KEGG_VALINE_LEUCINE_AND_ISOLEUCINE_DEGRADATION | 0.00206 | 0.02021 | -1.69144 | C2 |
| ZHOU_INFLAMMATORY_RESPONSE_LIVE_DN | 0.00018 | 0.00491 | -1.69066 | C2 |
| ZHOU_INFLAMMATORY_RESPONSE_FIMA_DN | 0.00018 | 0.00491 | -1.67877 | C2 |
| KEGG_PYRIMIDINE_METABOLISM | 0.00055 | 0.00910 | -1.67832 | C2 |

| | | | | |
|---|---|---|---|---|
| KEGG_PROPANOATE_METABOLISM | 0.00337 | 0.02772 | -1.67702 | C2 |
| REACTOME_PHASE_II_CONJUGATION | 0.00279 | 0.02507 | -1.67630 | C2 |
| REACTOME_PURINE_METABOLISM | 0.00336 | 0.02770 | -1.67336 | C2 |
| GAZDA_DIAMOND_BLACKFAN_ANEMIA_PROGENITOR_DN | 0.00111 | 0.01415 | -1.67156 | C2 |
| KEGG_NUCLEOTIDE_EXCISION_REPAIR | 0.00301 | 0.02603 | -1.64967 | C2 |
| MOOTHA_VOXPHOS | 0.00184 | 0.01880 | -1.64118 | C2 |
| TRAYNOR_RETT_SYNDROM_DN | 0.00683 | 0.04488 | -1.63971 | C2 |
| IVANOVA_HEMATOPOIESIS_LATE_PROGENITOR | 0.00017 | 0.00491 | -1.62837 | C2 |
| KIM_GERMINAL_CENTER_T_HELPER_UP | 0.00260 | 0.02380 | -1.62570 | C2 |
| REACTOME_NUCLEOTIDE_EXCISION_REPAIR | 0.00559 | 0.03969 | -1.59902 | C2 |
| KRIEG_KDM3A_TARGETS_NOT_HYPOXIA | 0.00073 | 0.01074 | -1.59135 | C2 |
| REACTOME_DNA_REPAIR | 0.00201 | 0.01987 | -1.59088 | C2 |
| KEGG_OXIDATIVE_PHOSPHORYLATION | 0.00420 | 0.03262 | -1.55061 | C2 |
| NIELSEN_GIST | 0.00583 | 0.04085 | -1.54216 | C2 |
| BOSCO_INTERFERON_INDUCED_ANTIVIRAL_MODULE | 0.00661 | 0.04398 | -1.53940 | C2 |
| SHEN_SMARCA2_TARGETS_UP | 0.00035 | 0.00718 | -1.53731 | C2 |
| REACTOME_OLFACTORY_SIGNALING_PATHWAY | 0.00727 | 0.04637 | -1.51523 | C2 |
| DAIRKEE_TERT_TARGETS_DN | 0.00587 | 0.04106 | -1.51047 | C2 |
| PLASARI_TGFB1_TARGETS_10HR_DN | 0.00182 | 0.01880 | -1.50374 | C2 |
| GRAESSMANN_RESPONSE_TO_MC_AND_SERUM_DEPRIVATION_UP | 0.00238 | 0.02236 | -1.48426 | C2 |
| IVANOVA_HEMATOPOIESIS_INTERMEDIATE_PROGENITOR | 0.00459 | 0.03501 | -1.47965 | C2 |
| KIM_MYC_AMPLIFICATION_TARGETS_UP | 0.00311 | 0.02665 | -1.46881 | C2 |
| ZWANG_DOWN_BY_2ND_EGF_PULSE | 0.00235 | 0.02223 | -1.45721 | C2 |
| GARY_CD5_TARGETS_DN | 0.00035 | 0.00718 | -1.45582 | C2 |
| HOLLMANN_APOPTOSIS_VIA_CD40_DN | 0.00199 | 0.01981 | -1.45263 | C2 |
| IVANOVA_HEMATOPOIESIS_EARLY_PROGENITOR | 0.00035 | 0.00718 | -1.45039 | C2 |

| | | | | |
|---|---|---|---|---|
| GRAESSMANN_APOPTOSIS_BY_SERUM_DEPRIVATION_UP | 0.00035 | 0.00718 | -1.44759 | C2 |
| TAKEDA_TARGETS_OF_NUP98_HOXA9_FUSION_10D_UP | 0.00479 | 0.03581 | -1.44660 | C2 |
| JOHNSTONE_PARVB_TARGETS_2_DN | 0.00089 | 0.01202 | -1.44533 | C2 |
| MANALO_HYPOXIA_DN | 0.00107 | 0.01415 | -1.43679 | C2 |
| MULLIGHAN_MLL_SIGNATURE_2_UP | 0.00088 | 0.01201 | -1.40956 | C2 |
| KAUFFMANN_DNA_REPAIR_GENES | 0.00486 | 0.03612 | -1.40524 | C2 |
| REACTOME_GENERIC_TRANSCRIPTION_PATHWAY | 0.00160 | 0.01766 | -1.39094 | C2 |
| DOUGLAS_BMI1_TARGETS_DN | 0.00286 | 0.02511 | -1.37669 | C2 |
| COLINA_TARGETS_OF_4EBP1_AND_4EBP2 | 0.00321 | 0.02726 | -1.36641 | C2 |
| STEIN_ESRRA_TARGETS_UP | 0.00250 | 0.02308 | -1.35620 | C2 |
| CHICAS_RB1_TARGETS_CONFLUENT | 0.00244 | 0.02264 | -1.31838 | C2 |
| SENESE_HDAC3_TARGETS_DN | 0.00299 | 0.02596 | -1.30385 | C2 |
| BACH2_01 | 0.00022 | 0.00123 | 2.02391 | C3 |
| ATF_01 | 0.00022 | 0.00123 | 2.00365 | C3 |
| BACH1_01 | 0.00022 | 0.00123 | 1.91461 | C3 |
| AP1_01 | 0.00022 | 0.00123 | 1.89068 | C3 |
| CREL_01 | 0.00022 | 0.00123 | 1.80675 | C3 |
| CREB_01 | 0.00022 | 0.00123 | 1.78187 | C3 |
| MYCMAX_01 | 0.00022 | 0.00123 | 1.74545 | C3 |
| WHN_B | 0.00022 | 0.00123 | 1.71762 | C3 |
| E2A_Q2 | 0.00022 | 0.00123 | 1.63008 | C3 |
| CACBINDINGPROTEIN_Q6 | 0.00022 | 0.00123 | 1.62430 | C3 |
| NFKB_C | 0.00022 | 0.00123 | 1.57477 | C3 |
| TATA_C | 0.00067 | 0.00260 | 1.57262 | C3 |
| EGR1_01 | 0.00045 | 0.00195 | 1.57223 | C3 |

| | | | | |
|---|---|---|---|---|
| ETS1_B | 0.00022 | 0.00123 | 1.55797 | C3 |
| XBP1_01 | 0.00328 | 0.00951 | 1.54089 | C3 |
| CGTSACG_PAX3_B | 0.00286 | 0.00857 | 1.53348 | C3 |
| TAL1ALPHAE47_01 | 0.00202 | 0.00659 | 1.49712 | C3 |
| CTACCTC_LET7A_LET7B_LET7C_LET7D_LET7E_LET7F_MIR98_LET7G_LET7I | 0.00023 | 0.00123 | 1.47711 | C3 |
| FOXO1_01 | 0.00465 | 0.01253 | 1.46751 | C3 |
| GFI1_01 | 0.00447 | 0.01209 | 1.43151 | C3 |
| PITX2_Q2 | 0.00829 | 0.01961 | 1.39842 | C3 |
| TAL1BETAITF2_01 | 0.00918 | 0.02116 | 1.38816 | C3 |
| CEBP_01 | 0.00918 | 0.02116 | 1.38620 | C3 |
| P53_DECAMER_Q2 | 0.00894 | 0.02079 | 1.38342 | C3 |
| ETS2_B | 0.01015 | 0.02240 | 1.35701 | C3 |
| TAL1BETAE47_01 | 0.01771 | 0.03423 | 1.34004 | C3 |
| OCT1_03 | 0.02456 | 0.04264 | 1.33957 | C3 |
| FOXO3_01 | 0.01737 | 0.03400 | 1.33916 | C3 |
| E47_01 | 0.01948 | 0.03675 | 1.32723 | C3 |
| FOXO4_02 | 0.02301 | 0.04096 | 1.31190 | C3 |
| E2F_02 | 0.02263 | 0.04072 | 1.30357 | C3 |
| YATGNWAAT_OCT_C | 0.02000 | 0.03718 | 1.29334 | C3 |
| USF_C | 0.02490 | 0.04291 | 1.29008 | C3 |
